# Genomic organization of the autonomous regulatory domain of *eyeless* locus in *Drosophila melanogaster*

**DOI:** 10.1101/2021.04.15.440022

**Authors:** Shreekant Verma, Rashmi U Pathak, Rakesh K Mishra

## Abstract

In *Drosophila*, expression of *eyeless* (*ey*) gene is restricted to the developing eyes and central nervous system. However, the flanking genes, *myoglianin* (*myo*), and bent (*bt*) have different temporal and spatial expression patterns as compared to the *ey*. How distinct regulation of *ey* is maintained is mostly unknown. Earlier, we have identified a boundary element intervening *myo* and *ey* genes (ME boundary) that prevents the crosstalk between the *cis*-regulatory elements of *myo* and *ey* genes. In the present study, we further searched for the *cis*-elements that define the domain of *ey* and maintain its expression pattern. We identify another boundary element between *ey* and *bt*, the EB boundary. The EB boundary separates the regulatory landscapes of *ey* and *bt* genes. The two boundaries, ME and EB, show a long-range interaction as well as interact with the nuclear architecture. This suggests functional autonomy of the *ey* locus and its insulation from differentially regulated flanking regions. We also identify a new Polycomb Response Element, the *ey*-PRE, within the *ey* domain. The expression state of the *ey* gene, once established during early development is likely to be maintained with the help of *ey*- PRE. Our study proposes a general regulatory mechanism by which a gene can be maintained in a functionally independent chromatin domain in gene-rich euchromatin.

## Introduction

The *Drosophila eyeless* gene (*ey*), a *Pax*-6 homolog, is an essential regulatory gene required for development of the eye (Quiring *et al*. 1994). It is one of the many evolutionarily conserved genes that are involved in retinal determination. Molecular genetic studies have shown that *ey* along with other genes, viz., *eyes absent* (*eya*), *twin of eyeless* (*toy*), *sine oculis* (*so*), and *dachshund* (*dac*), is involved in retinal determination network (Desplan 1997; Halder *et al*. 1998; Pichaud AND DESPLAN 2001). Although the genetic interactions of *ey* with these genes are mostly known, spatial and temporal transcriptional regulation of the *ey* is less well understood.

The *ey* gene, present on the fourth chromosome of *Drosophila*, is flanked closely by *myoglianin* (*myo*) gene upstream, and *bent* (*bt*) gene downstream. The three genes show a very distinct spatiotemporal pattern of expression in the fly. The expression of *ey* is restricted to eye disc primordia and central nervous system (CNS) during embryogenesis. Later during larval stages it is expressed in eye imaginal discs and CNS where it continues to express in adults (Quiring *et al*. 1994; Hauck *et al*. 1999; Adachi *et al*. 2003; Chintapalli *et al*. 2007). Unlike *ey,* the *myo* gene that codes for a TGF-β superfamily protein, has a high-level of maternal β transcript deposition in the early embryo. Later, it expresses in the somatic, visceral, and heart musculature where its expression persists throughout embryogenesis. In third instar larvae, *myo* expression is restricted to brain and glial cells of the ventral nerve cord (Lo AND FRASCH 1999; Chintapalli *et al*. 2007). Likewise, the *bt* gene encodes for a titin superfamily muscle protein ‘Projectin’. During development, the *bt* transcripts first appear in mid-embryo stage and increase steadily till adult stage (Fyrberg *et al*. 1992; Maroto *et al*. 1992). The expression of *bt* has been found in the embryonic/larval muscles system and all types of muscles in pupae and adults (Ayme-southgate *et al*. 1991; Frise *et al*. 2010).

Transcriptional regulation of the *ey* at various developmental stages by multiple *ey* enhancers has been reported in two earlier studies (Hauck *et al*. 1999; Adachi *et al*. 2003). An initial study has identified a 212-bp eye-specific enhancer at 3’ end of the second intron that is essential for expression of *ey* in larval eye-disc primordia (Hauck *et al*. 1999). A later study has described a 5 Kb upstream region and a 3.6

Kb second intronic fragment that act synergistically to specify the *ey* specific expression pattern of lacZ reporter construct in developing CNS (Adachi *et al*. 2003). The targeted expression of *ey* induces ectopic eyes in *Drosophila* (Halder *et al*. 1998). This suggests that active repression of the *ey* is crucial in other tissues, wherever it is not required. However, what maintains the repressed status of *ey* in such tissues, is not known. The distinct spatiotemporal expression of the *ey* calls for the presence of other *cis*-regulatory elements like chromatin domain boundary elements (CBEs) or insulators, and also a memory element like Polycomb response elements (PREs). Such *cis*-elements at the *ey* locus are required to prevent the cross-talk among the regulatory elements of the three genes and for maintenance of tissue-specific expression, as is the case at several well characterised loci like bithorax complex in *Drosophila* (Maeda AND KARCH 2006; Mihaly *et al*. 2006).

Boundary elements are established regulatory feature of most eukaryotic genomes. They insulate a gene from the influence of neighbouring chromatin and actively subdivide the genome into chromatin domains of independent gene activity. The regulatory function of these elements is dependent on their association with several boundary interacting proteins including BEAF-32, CTCF, Su(Hw), Zw5, GAGA factor (GAF or Trl) and CP190 (Geyer *et al*. 1986; Kellum AND SCHEDL 1991; Gerasimova AND CORCES 2001; West *et al*. 2002; Pai *et al*. 2004; Gaszner AND FELSENFELD 2006; Valenzuela AND KAMAKAKA 2006). On the other hand, PREs, also known as <u>C</u>ellular <u>M</u>emory <u>M</u>odules (CMMs) are specific DNA elements, recruitment sites for Polycomb group and trithorax group (PcG/trxG) of proteins, that are required to maintain the transcriptional states of the target genes (Sigrist AND PIRROTTA 1997; Cavalli AND PARO 1998). The PcG proteins are required for stable silencing, whereas trxG proteins promote activation of target genes. Many PREs studied in *Drosophila* suggest their role in maintenance of repressed states (Kassis AND BROWN 2013). One such example is *iab7*-PRE in the *Drosophila* bithorax complex, The *iab7*-PRE helps in appropriate maintenance of *Abd-B* gene expression pattern in parasegement 12 (Mishra *et al*. 2001). Studies based on techniques such as FISH and chromosome conformation capture (3C) assay, support interactions among CBEs and other regulatory elements like promoters, enhancers and silencers. This leads to the formation of chromatin domains with autonomous regulatory function (Dekker *et al*. 2002; Bantignies *et al*. 2003; Ronshaugen AND LEVINE 2004; Lanzuolo *et al*. 2007; Sexton *et al*. 2009; Hou *et al*. 2012; Zhang *et al*. 2013).

These higher order organization of chromatin interaction are known as topological associated domains (TADs). (Hou *et al*. 2012; Sexton *et al*. 2012; Zhang *et al*. 2013; Cubenas-potts *et al*. 2017). In addition, anchoring of regulatory elements to nuclear architecture also helps in compartmentalization of a locus into functionally independent domain (Pathak *et al*. 2007).

In the present study, we have analyzed the *ey* locus to identify *cis-* regulatory elements that define its precise and distinct expression pattern. Earlier we have identified a CBE upstream of *ey* that separates regulatory domains of *myo* and *ey* genes (Sultana *et al*. 2011). Here we have identified a CBE between *ey* and *bt*, two closely spaced but distinctly expressed genes. We refer this element as EB boundary and functionally characterize it in the transgenic context. Additionally, we have identified a PRE (*ey*-PRE) that functions as a repressor to maintain the expression pattern of *ey*. Furthermore, to understand the mechanism of *ey* regulation by EB, *ey-*PRE, and previously identified ME boundary, we have explored their long-range interactions in the context of nuclear architecture. Our findings reveal two new regulatory elements in this important locus, and the structural framework of this genomic locus based on the higher-order chromatin organization of regulatory elements. Such organization may have a general implication in higher eukaryotes.

## Results

### Identification of potential CBEs that separate ey and bt loci

Previously, we have identified a CBE (the ME boundary) in the ∼1.6 Kb intergenic region between *myo* and *ey* genes, endorsing the rationale that CBEs are required to prevent crosstalk between the regulatory elements of closely spaced but differently expressed genes (Sultana *et al*. 2011). Likewise, *ey* and the downstream gene *bt,* have a disparate pattern of expression (Ayme-southgate *et al*. 1991). Therefore, we expected a CBE to be present in the 3.2 Kb intergenic region to block crosstalk between the adjacent regulatory domains. To identify a potential CBE between *ey* and *bt*, we used cdBEST search tool which has been successfully utilized to identify several new CBEs in *Drosophila* and other insects like *Anopheles gambie* (Srinivasan AND MISHRA 2012; Ahanger *et al*. 2013). CdBEST predicts CBEs based on the presence of clusters of binding motifs of many known boundary proteins like BEAF-32, CTCF, GAF, CP190, and *Zw5,* in the queried sequence.

CdBEST analysis for the ∼30 Kb *ey* locus predicted two CBEs, one being the previously reported ME boundary and a new putative CBE in the seventh intron of *ey* gene (INT7) (Figure 1). No CBE is predicted in the intergenic region between *ey* and *bt,* although the region carried multiple binding motifs of BEAF-32, CTCF, and GAF. Therefore, we analyzed the data available for *in-vivo* binding of boundary proteins in this region using embryonic (0-12 h) genome-wide ChIP-chip datasets from the modENCODE project (http://www.modencode.org/) (Negre *et al*. 2010). We observed a significantly high occupancy of CTCF and CP190 in a ∼1.2 Kb region within INT7. However, a prominent binding of both of these proteins is also observed in the ∼2.2 Kb region between *ey* and *bt* (EB). Particularly, CTCF binding at EB occurs in two separate regions, one in a ∼0.8 Kb 3’ UTR region of *ey* (named as EB- u), and the other in adjacent ∼1.1 Kb intergenic region (named as EB-i). EB-i additionally has binding of BEAF-32. We did not find a significant binding of GAF in these regions (Figure 1).

**Figure 1.**
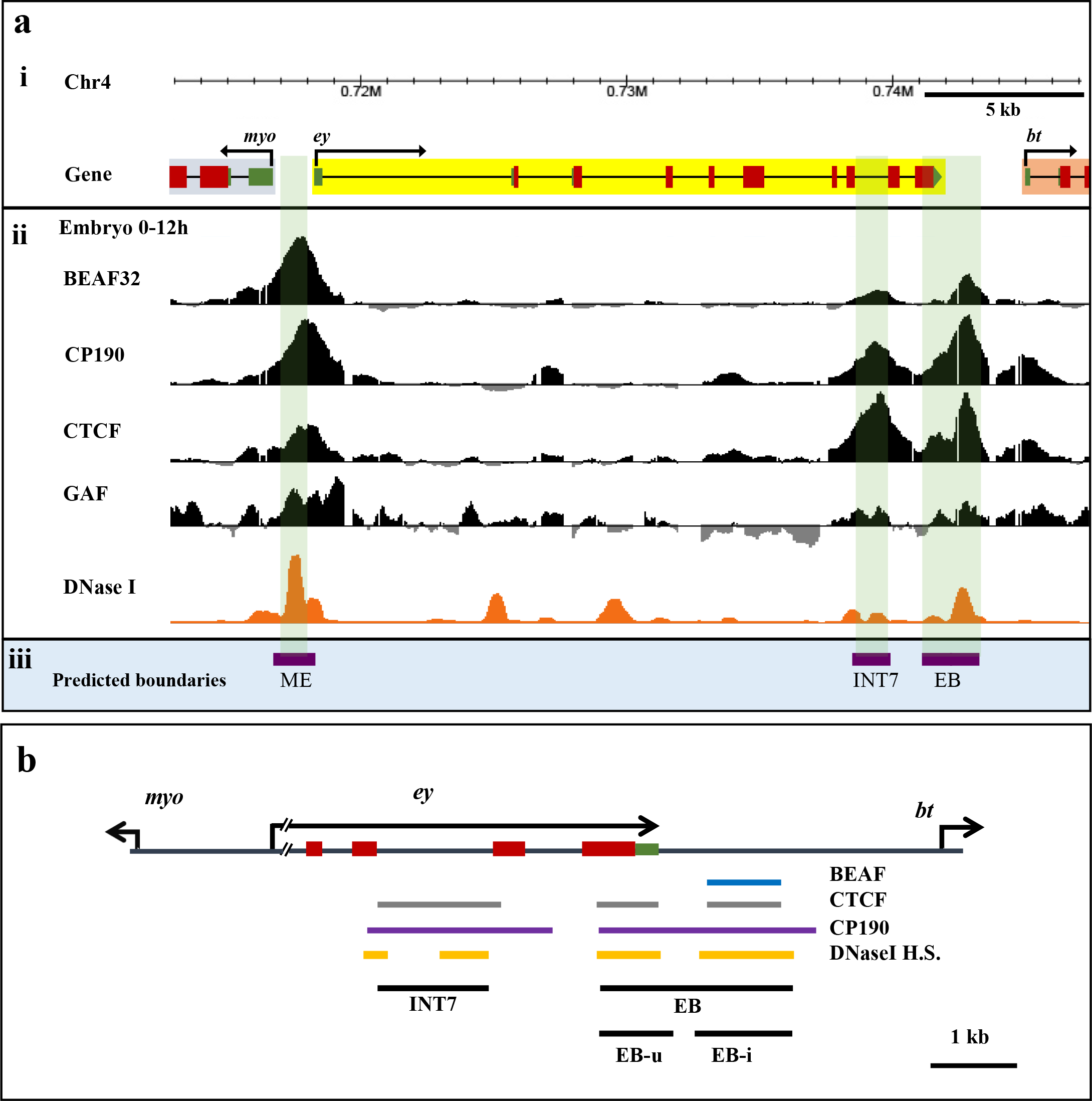
Prediction of CBE at the *ey* locus. **(a) (i)** *ey* locus on the fourth chromosome of *Drosophila melanogaster* showing *myo* and *bt* genes (genome version dm3). **(ii)** Mapping of boundary proteins binding sites and DHS. ChIP-chip data for binding of BEAF-32, CP190, CTCF, and GAF in 0-12 h embryos is used from Negre et al. (Negre *et al*. 2010). For mapping of DHS, data from Thomas et al. for ∼6 h embryo is used (Thomas *et al*. 2011). **(iii)** Prediction of CBEs. Previously reported ME boundary between *ey* and *myo* and a new CBE at INT7 in the seventh intronic region of *ey* is predicted by cdBEST. Binding of boundary proteins and DHS coincide with ME, INT7 (∼1.2 Kb) and EB (∼2.2 Kb) genomic regions (highlighted) although EB is not predicted as a CBE. **(b)** Map showing binding of boundary proteins (BEAF-32, CTCF and CP190) and DHS in INT7 and EB region that are tested as enhancer blockers in the present study. EB is further divided into EB-u and EB-i based on CTCF binding and presence of DHS.

As the existence of DNaseI hypersensitive sites (DHS) are inherent feature of regulatory elements, including several known CBEs, we next examined the presence of DHS in INT7 and EB using the DHS dataset of embryonic stage 9 (6 h embryo) available at UCSC genome browser (http://genome.ucsc.edu/) (Thomas *et al*. 2011). Both INT7 and EB contained a prominent DHS (Figure 1a and b). These observations altogether indicate potential CBE function of INT7 and EB (Figure 1b, Supplementary Figure 1, Supplementary Table 1).

### EB functions as an enhancer blocker in the eye

To test the boundary activity of INT7 and EB, we generated transgenic flies carrying test fragments introduced between the *white* enhancer and *mini*-*white* reporter gene in enhancer blocker assay construct (pRW+) (Figure 2a) (Hagstrom *et al*. 1996). The level of eye pigmentation in adult transgenic flies is a responsive indicator of the amount of *mini-white* transcription. In case a CBE placed between the enhancer and *mini-white* acts as an enhancer blocker, it would reduce the level of *mini*- *white* expression that can be scored as light eye colour in adult flies. However, a random DNA fragment in the same position would have no effect (Hagstrom *et al*. 1996). The test fragments in the assay vector are flanked by *lox*P sites so that it can be flipped out using the *cre-lox*P system to confirm the enhancer blocker effect and rule out any position effect.

**Figure 2.**
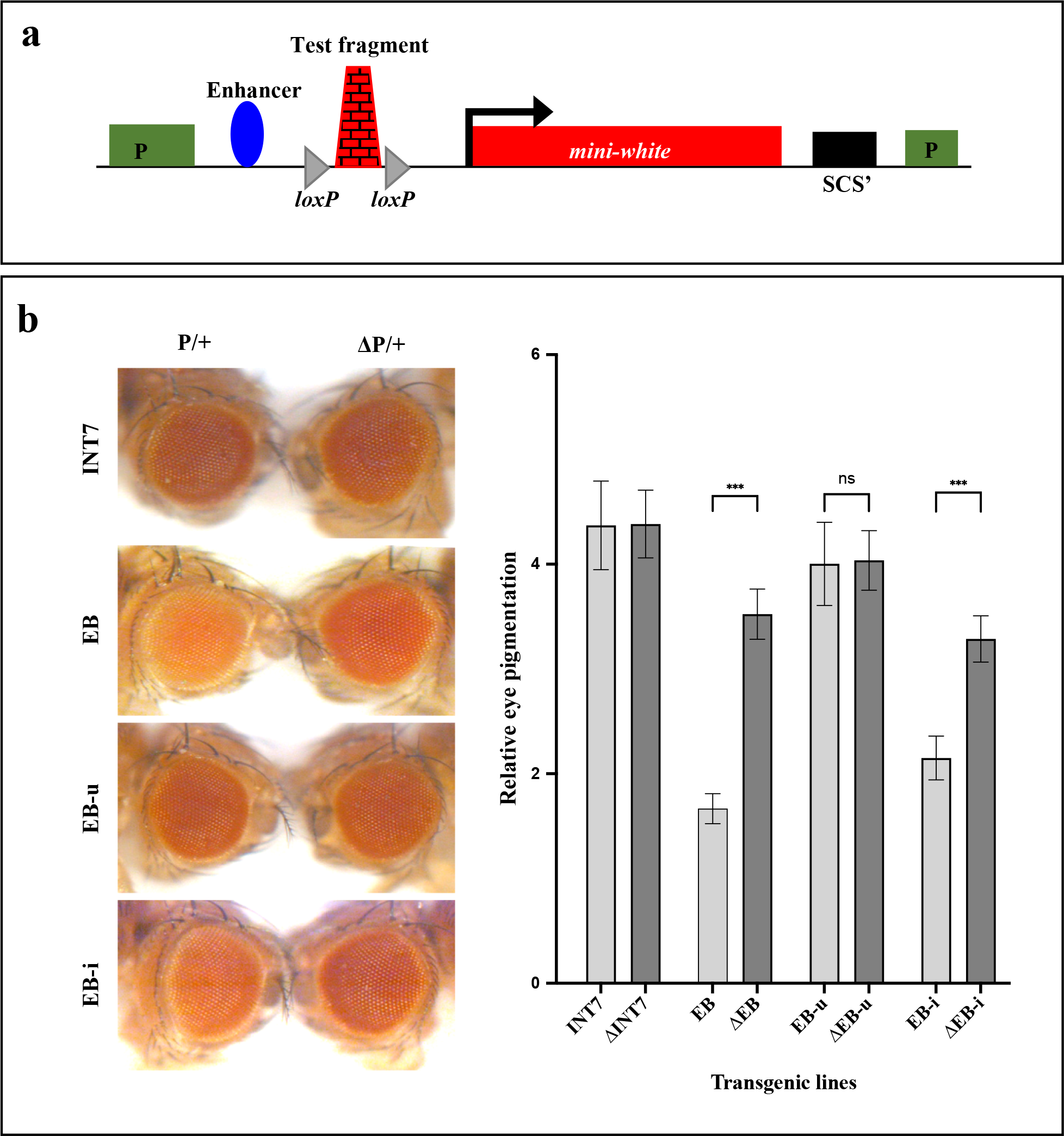
EB functions as Enhancer blocker in adult eye. **(a)** Enhancer blocker assay construct, pRW^+^. The test fragment flanked with *loxP* sites is inserted between the enhancer and promoter of *mini-white*. **(b)** Enhancer blocker assay in the eye. Eye colour comparisons of one of the transgenic lines from each fragment tested are shown (Supplementary Table 2). In each panel, the eye on the left is from a transgenic fly that contains the test fragment in heterozygous (P/+), whereas the eye on the right is from a fly of the same line after the fragment is flipped out (Δ +). P/ INT7 region does not show enhancer blocker activity while EB shows a weak enhancer blocker activity. EB-u and EB-i, were tested separately, and only EB-i shows activity similar to EB suggesting that the boundary function of EB resides in EB-i. The graph represents mean and standard deviation of relative pigment value in the eyes from three experiments with 10 heads per experiment. The mean, standard deviations and the significance of differences were analyzed by one-way ANOVA where α=0.05, p<0.001(_***_) and ns=non significant.

All the transgenic fly lines generated for INT7 (∼10 lines) had a high level of eye pigmentation (red to bright red eye colour) in heterozygous condition. Moreover, the removal of INT7 in 4 initial lines (P) did not change in the level of eye pigment in their flipped-outs (Δ ) indicating that INT7 lacked an enhancer-blocker activity (Figure 2b, Supplementary Table 2). In contrast, 7 transgenic lines carrying EB fragment displayed varying degrees of light red eye colour in heterozygous condition and 4 of them turned darker when EB was flipped out. We found a mild increase in eye colour in 3 EB flipped-out lines and a moderate (∼ 2 fold) increase in eye pigment level in 1 EB flipped out line (Figure 2b, Supplementary Table 2). Homozygous (P/P) transgenic lines mostly showed a high level of eye pigmentation (dark red eye colour) in both the INT7 and EB transgenes. These results suggested that the EB acts as a weak enhancer blocker in the eye and the flipped-out lines confirmed that light eye colour is due to the boundary function of EB and not a position effect.

As explained in the earlier section, EB can be split into two fragments based on CTCF binding sites as EB-u and EB-i. The binding of BEAF-32, however, is detected only in the EB-i region. Both of these regions contain distinct DHS, which is more prominent in the EB-i (Figure 1b). We further tested the enhancer blocking activity of both the fragments separately, to identify whether the boundary function lies in one of these regions or the entire EB sequence is required for the activity. All 7 EB-u transgenic lines tested had dark red eye colour and did not exhibit enhancer-blocker effect as no change in eye colour was observed in flipped-out lines. In contrast, EB-i transgenic lines showed light eye colour and 5 out of 7 lines showed a mild (∼1.5 fold) increase in eye colour after deletion of EB-i, suggesting that enhancer blocker activity of EB mainly resides in EB-i (Figure 2b, Supplementary Table 2).

In summary, the presence of DHS with significant boundary proteins occupancy at INT7 and EB suggested their boundary features, but only EB displayed enhancer blocker effect in the adult eye. As the CBE features were observed using datasets from embryonic developmental stages, we reasoned that these regions might be functional as a boundary mostly during early development. Therefore, we tested the boundary activity of INT7 and EB in the embryo.

### EB functions as an enhancer blocker in embryo

We used a P-element based CfhL assay vector to test boundary activity in developing embryos (Figure 3a) (Hagstrom *et al*. 1996). The construct carries two reporter genes *mini-white* and *hsp70/lacZ.* Two *ftz* enhancers, an upstream enhancer (UPS) active during early development and a late neurogenic enhancer (NE), drive the *hsp70/lacZ* in the embryo. The level of eye pigmentation and X-gal (lacZ) staining in the embryo both indicates the expression level of *mini-white* and *lacZ*, respectively. A test DNA that acts as an enhancer blocker when inserted between the *lacZ* promoter and the enhancers would cause a reduced level of *lacZ* expression. The test element can be flipped out from the transgenes to confirm the enhancer-blocker activity and rule out any position effect. In addition, the *mini-white* expression level in the adult eye can be compared to rule out a repressor effect of the test elements (Hagstrom *et al*. 1996; Sultana *et al*. 2011).

**Figure 3.**
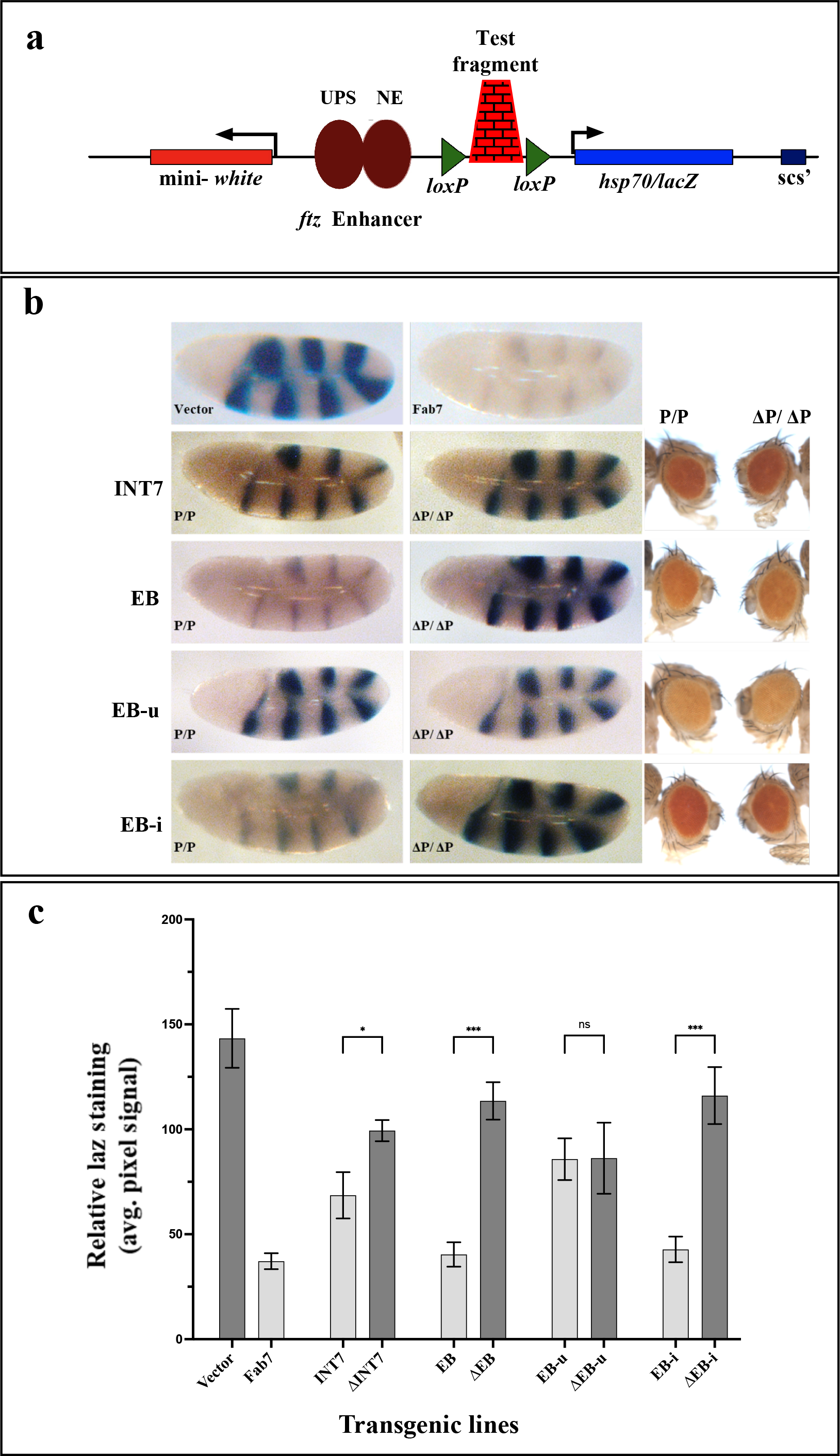
EB functions as an Enhancer blocker in the embryo. **(a)** Enhancer blocker assay construct, pCfhL. If a test DNA, inserted between the *ftz* enhancers and *hsp70*/*lacZ* gene, functions as a boundary, it prevents the enhancer from driving the reporter gene, thereby, giving a low intensity of lacZ staining, which is enhanced upon flipping out of the test DNA. **(b**) LacZ staining in embryos. The panel compares the *ftz* UPS enhancer mediated lacZ expression in homozygous transgenic embryos during early development (P/P) and embryos obtained from the same line afterflipping out of the transgene (Δ ΔP). Transgenic lines of INT7, EB, EB-u, and EB-iP/ were tested (Supplementary Table 3). Transgenic lines with pCfhL construct alone and with Fab7 boundary were used as negative and positive controls, respectively. INT7 shows a weak enhancer blocker effect while EB shows a significant enhancer blocking activity. No significant change in eye colour in flipped out lines was observed (Panel on the right). Further, the boundary activity of EB resides in EB-i, whereas EB-u does not show enhancer blocker effect. **(c)** Quantification LacZ staining. Using ImageJ tool , the mean pixels value of a fixed area of lacZ stained regions of each genotype were quantitated. For each genotype, 3-5 embryos were taken which represented prevalent staining pattern of the group. The relative lacZ staining, standard deviations and the significance of differences in the graph were analyzed by one-way ANOVA where α=0.05, p<0.01(_*_), p<0.001(_***_) and ns=non significant.

In EB transgenic lines, we observed significantly reduced lacZ staining in both early and late embryonic stages suggesting that EB prevented both the enhancers from acting on *lacZ* (Figure 3b-c and Supplementary Figure 2). EB in homozygotic condition displayed a strong boundary activity which is similar to the known *Fab7* boundary (Hagstrom *et al*. 1996). In 2 out of 3 EB flipped out version of the transgenic lines, an intense lacZ staining was restored (comparable to empty vector). At the same time, the eye colour remained the same, suggesting that EB functions as an enhancer blocker but not a repressor (Figure 3b-c, Supplementary Figure 2 and Supplementary Table 3). Interestingly, INT7 transgenes also displayed a weak enhancer blocker activity in the embryo. Two out of the 3 transgenic lines of INT7 showed a moderate increase in the lacZ staining in both early and late stages embryos after removal of the test elements (Figure 3b-c, Supplementary Table 3, Supplementary Figure 2). When EB-u and EB-i fragments were tested separately, EB-u (0/3) did not exhibit any enhancer blocker activity in the embryo as an intense lacZ staining comparable to empty vector was observed. The EB-i region, on the other hand, showed a strong enhancer blocker activity, comparable to EB. Like EB, flipped out EB-i (3/5) lines showed a significant increase in the lacZ staining with no change in eye colour. These observations indicated that mainly the EB-i, which carries BEAF-32 binding sites harbours the boundary activity in this region (Figure 3b-c, Supplementary Figure 2 and Supplementary Table 3). Altogether these results suggest that EB and INT7 are active boundaries during embryonic development. Further, a weak enhancer blocker activity of INT7 in the embryo suggest its assistive function to EB (a strong boundary) in demarcating the *ey* locus.

### Identification of a putative PRE associated with ey locus

It is known that multiple enhancers positively regulate the *ey* gene in developing eye and CNS (Hauck *et al*. 1999; Adachi *et al*. 2003). However, the regulatory elements that maintain the repressed state of *ey* in other tissues are not known. In *Drosophila,* PREs are involved in establishing and keeping the repressed state of genes (Mishra *et al*. 2001; Americo *et al*. 2002). We therefore searched for putative PRE in *ey* locus using a Perl-based PRE mapper tool (Srinivasan AND MISHRA 2020). This tool uses a motif cluster search of known DNA binding PcG recruiters like Pho, GAF, Dsp1, Sp1, and Cg. PRE mapper predicted a ∼1.6 Kb putative PRE region upstream of the exon 1 of *ey.* We named this element as *ey*-PRE, and observed that it has clusters of Sp1 (3), Cg (3), Pho (1), and GAF (1) binding sites in an internal core region of ∼800 bp. Interestingly, the *ey*-PRE also contained a second *ey* promoter (P2) region as predicted by eukaryotic promoter database (Dreos *et al*. 2015) (Figure 4).

**Figure 4.**
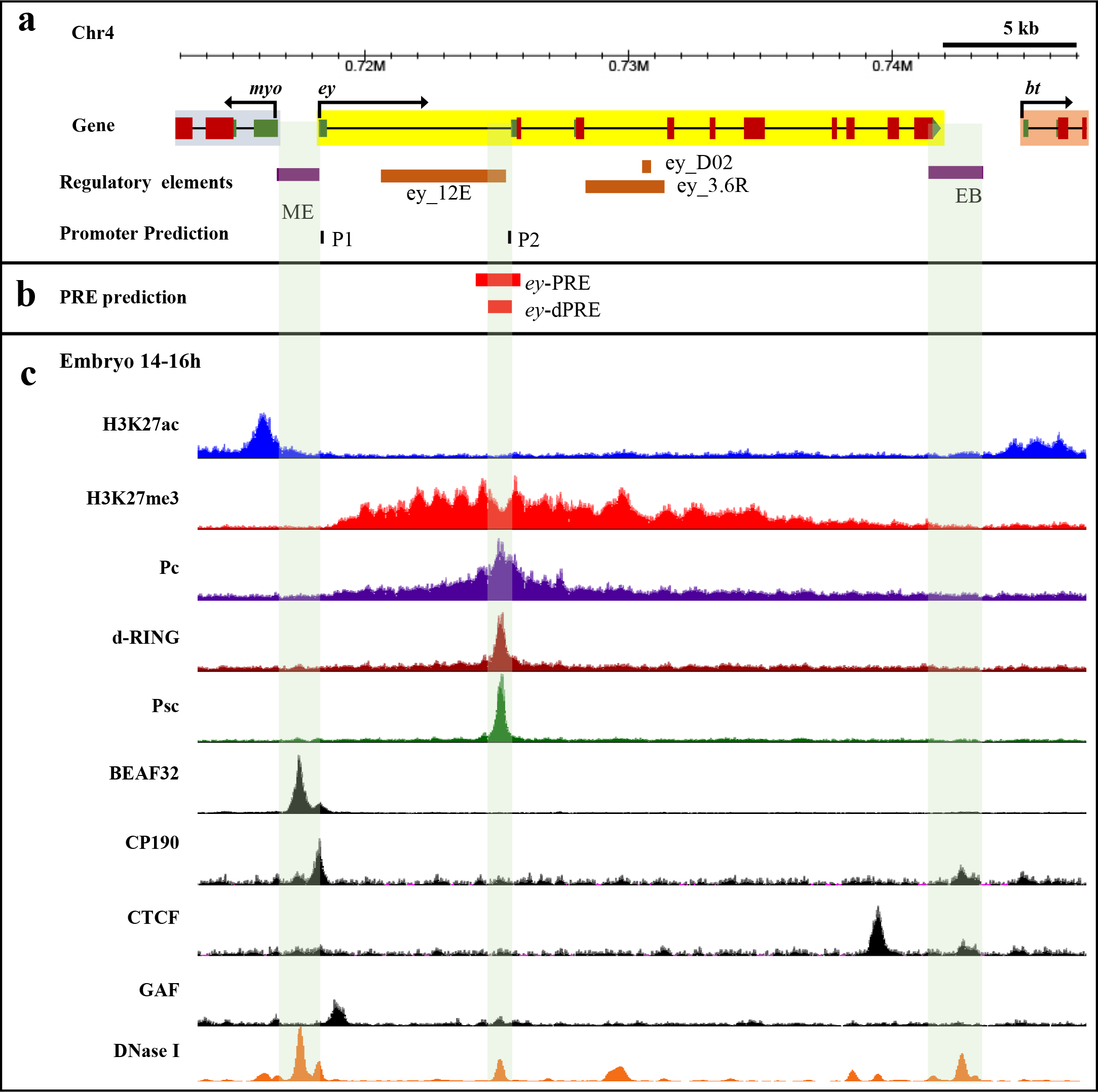
PRE prediction in *ey* locus. **(a)** *Cis*-regulatory elements at *ey* locus. Specific enhancers of *ey* (shown in orange), identified in previous studies are present in 5’-UTR (*ey_12E)* and in the second intron (*ey_3.6R* and *ey_D02)* (Hauck *et al*. 1999; Adachi *et al*. 2003). The two CBEs, ME and EB, demarcate the *ey* locus. Two promoters, P1 and P2 are predicted at the locus by eukaryotic promoter database (Dreos *et al*. 2015). **(b)** A ∼1.6Kb *ey-*PRE and a core 831bp *ey-*dPRE in the *ey* locus is predicted by PRE mapper. The *ey* predicted promoter P2, *also* coincides with *ey-*PRE. **(c)** Histone modification profiles, binding sites of PcG and boundary proteins, and DHS at *ey* locus in the 14-16 h embryo. ChIP-seq data from the modENCODE project (Mod *et al*. 2010) and DHS data of stage 9 (∼6h) embryo from Thomas et al. 2011 (Thomas *et al*. 2011) have been used at the UCSC browser to generate the map. The *ey* locus has a repressed H3K27me3 domain, whereas neighbouring genes *myo* and *bt,* both have active H3K27ac marks at their promoters. Binding of Pc spreads over the entire *ey* gene while dRING, and Psc, along with Pc, show a sharp peak upstream to the first exon and align perfectly with the predicted *ey-*PRE. The core of *ey*-PRE maps to DHS and is named as *ey-*dPRE. Boundary proteins BEAF-32, CP190, and CTCF bind at both ME and EB that appear to form a boundary to the repressed domain of *ey*.

To assess whether the *ey-*PRE is a target of PcG proteins, we looked into the presence of repressive histone modification (H3K27me3) and binding of PcG proteins (PRC1 members - Pc, Psc, and dRing) (Figure 4c). We used data for 14-16 h embryo as this was the only stage for which a ChIP-seq data for all the queried proteins and histone modification was available at the modENCODE database (Mod *et al*. 2010). Binding of boundary proteins was somewhat comparable but not identical to previously used 0-12 h ChIP-chip data (Figure 1a). We observed a H3K27me3 domain with significant enrichment of Pc in *ey* locus. Interestingly, the repressive H3K27me3 modification is primarily confined to *ey* locus only, whereas flanking regions toward *myo* and *bt* containactive histone modification of H3K27ac. At the same time, boundary proteins namely BEAF-32, CP190, and CTCF (Chip-seq data from modENCODE project) are present at both, ME and EB boundaries that demarcate the borders of the repressive domain of *ey* (Figure 1a and 4c). Presence of a Pc-enriched repressive domain restricted to *ey* locus indicates that a repressive *cis*-acting element is very likely to be present within the locus. Additionally, binding of PcG proteins Pc, dRING, and Psc in a significantly high amount as a sharp peak at putative *ey-*PRE further strengthens the idea. The ∼800 bp internal core region also contained a DHS. With such strong indications for the presence of a functional PRE at the *ey* locus, we decided to test both *ey*-PRE and a smaller fragment of core DHS region, *ey*-dPRE, for PRE activity (Figure 4b, Supplementary Table 1).

### ey-PRE functions as a PRE

To determine the PRE potential of *ey*-PRE, we assessed the repressor activity of full-length *ey*-PRE (∼1.6Kb) and *ey*-dPRE (831 bp) in the adult eye using a previously described pCasper vector based transgene assay (Figure 5a). In this assay, the presence of a PRE upstream of the *mini-white* gene causes its repression resulting in reduced and variegating eye colour (Vasanthi *et al*. 2013). We observed both repression and variegation of *mini*-*white* in the *ey-*PREs transgenic lines. Of the *ey*-PRE transgenic lines, 52% (16 out of 31) lines showed lighter eye colour, and 55% (18 out of 31) lines showed variegation. In the case of *ey*-dPRE, 50% (13 out of 26) lines displayed lighter eye colour while 21% (4 out of 19) lines showed variegation (Figure 5b, Table 1, and Supplementary Table 4). Further, to confirm that the repressive activity of *ey-*PRE and *ey*-dPRE is not a position effect, we excised out the PREs. Excision of *ey*-PRE/*ey*-dPRE led to the de-repression of *mini-white* in both sets of transgenic lines (Figure 5b and Table 1). Specifically, 59% and 47% of *ey*-PRE and *ey*-dPRE lines, respectively, showed an increase in the eye colour after flipping out of the transgene. Interestingly, we also observed a loss of variegation in the flipped-out lines of both the transgenes confirming the repressor function of *ey*-PRE/*ey*-dPRE.

**Figure 5.**
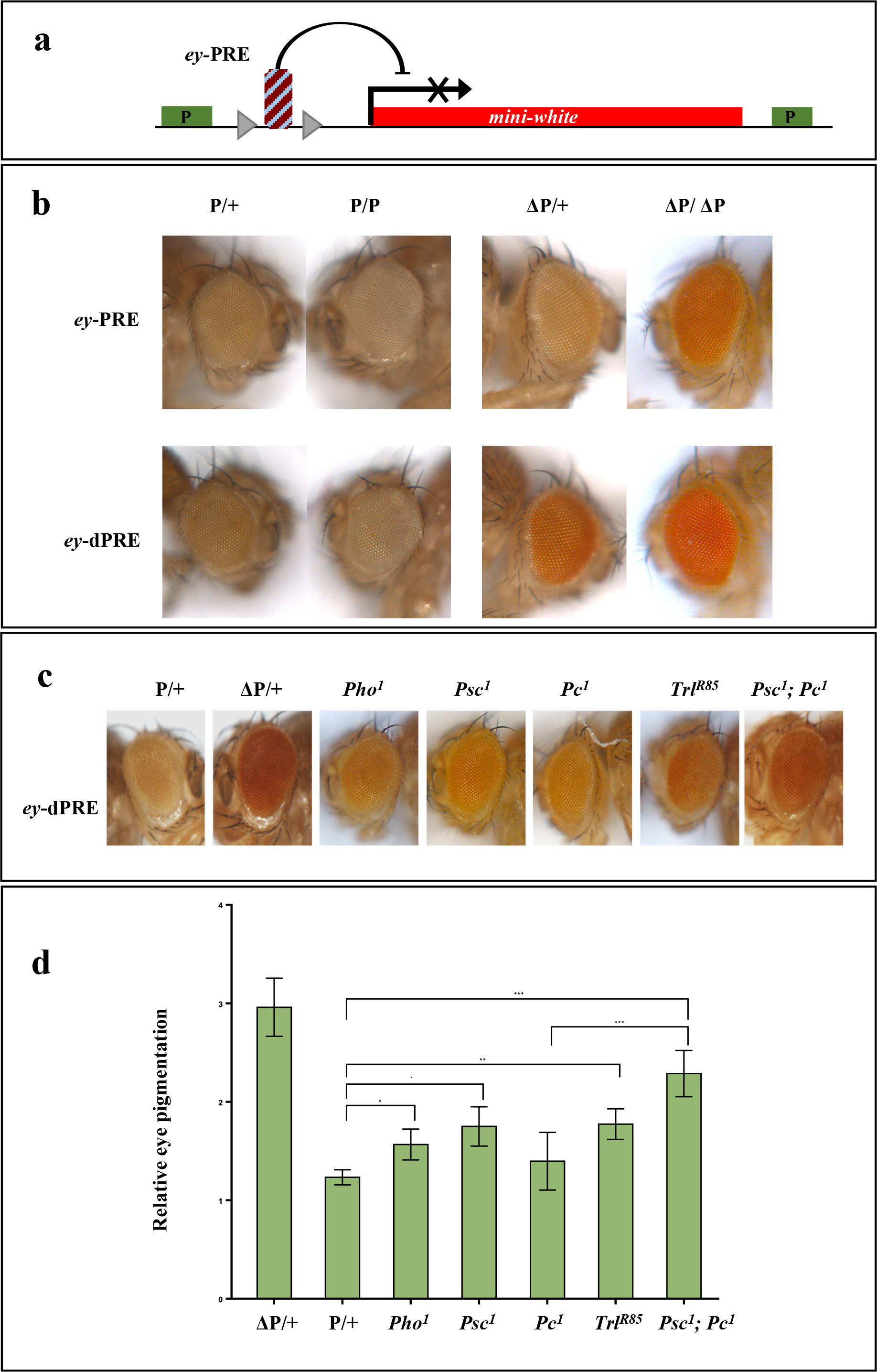
Functional validation of *ey*-PRE. **(a)** A pCaSpeR based PRE assay vector. The test DNA is inserted upstream to the *mini-white* gene in pCaSpeR vector. The test DNA is flanked by *loxP* sites. If the test DNA functions as a PRE, it would repress *mini-white*. The test fragments can be removed with the help of *loxP* to confirm the PRE mediated repression activity and rule out any position effect of the genomic environment. **(b)** *ey*-PREs function as a repressor and show strong PSS. Both, *ey*-PRE (∼1.6Kb) and *ey*-dPRE (831bp) exhibit strong repression of *mini-white* in homozygous flies (P/P) compared to their heterozygous counterparts (P/+). The flipped-out versions of the same (Δ + and Δ Δ show de-repression of *mini-white*. More than 50% of *ey-*PREs lines show strong repression of *mini-white* displayed as very light and variegating eye colour. The smaller version, *ey-* dPRE also shows repression activity to a similar extent suggesting that smaller fragment is sufficient for the PRE activity (see Table1 and Supplementary Table 4). **(c)** Effect of PcG mutations on *ey*-PRE. All eyes are from flies heterozygous for *ey-*PRE and PcG mutations. The graph represents quantitation of the eye colour in the heterozygous condition in wild type and different PcG proteins mutation background. The mean of relative eye pigment value (in triplicate with 10 fly heads in each replicate), standard deviations and the significance of differences were analyzed by one-way ANOVA where α=0.1, p=0.011(_*_), p=0.008(_**_), p<0.001(_***_) and ns=non significant. PcG mutations, *Pc^1^* and *Pho^1^* both show a mild de-repression effect while *Psc^1^* attenuates the *ey*-dPRE function significantly (see Table 2). Mutation in GAF (*Trl^R85^*) also attenuates the PRE. A double PcG mutation of *Pc^1^* and *Psc^1^* shows additive de-repression.

**Table 1:** Repressor activity of *ey-*PRE and *ey-*dPRE transgenes

|  | **ey-PRE** | **ey-dPRE** |
| --- | --- | --- |
| Total number of lines | 31 | 26 |
| % lines variegating | 58 (18/31) | 21 (4/19) |
| % lines with light eye colour | 52 (16/31) | 50 (13/26) |
| % lines showing PSS | 32 (7/22) | 36 (5/14) |
| % lines showing increase in eye colour in flipped-outs | 59 (16/27) | 47(8/17) |

**Table 2:** Effect of PcG mutations on *ey-*PRE mediated repression

| **ey-PRE lines** | **PcG Mutation** | **Effect** | **Strength** |
| --- | --- | --- | --- |
| **13.2** | *Pc*<sup>1</sup> | Derepression | + |
|  | *Psc*<sup>1</sup> | Derepression | ++ |
|  | *Pho*<sup>1</sup> | Derepression | + |
|  | *Trl*<sup>R85</sup> | Derepression | ++ |
|  | *Pc*<sup>1</sup>; *Psc*<sup>1</sup> | Derepression | +++ |
| **76.1** | *Pc*<sup>1</sup> | No effect |  |
|  | *Psc*<sup>1</sup> | Derepression | ++ |
|  | *Pho*<sup>1</sup> | No effect |  |
|  | *Trl*<sup>R85</sup> | Derepression | ++ |
|  | *Pc*<sup>1</sup>; *Psc*<sup>1</sup> | Derepression | ++ |
| **100.2** | *Pc*<sup>1</sup> | Derepression | + |
|  | *Psc*<sup>1</sup> | Derepression | ++ |
|  | *Pho*<sup>1</sup> | Derepression | + |
|  | *Trl*<sup>R85</sup> | Derepression | ++ |
|  | *Pc*<sup>1</sup>; *Psc*<sup>1</sup> |  |  |
| **88.1** | *Pc*<sup>1</sup> | Derepression | + |
|  | *Psc*<sup>1</sup> | Derepression | ++ |
|  | *Pho*<sup>1</sup> | Derepression | + |
|  | *Trl*<sup>R85</sup> | Derepression | ++ |
|  | *Pc*<sup>1</sup>; *Psc*<sup>1</sup> | Derepression | +++ |
| **10.2** | *Pc*<sup>1</sup> | No effect |  |
|  | *Psc*<sup>1</sup> | Derepression | ++ |
|  | *Pho*<sup>1</sup> | Derepression | + |
|  | *Trl*<sup>R85</sup> | Derepression | ++ |
|  | *Pc*<sup>1</sup>; *Psc*<sup>1</sup> | Derepression | +++ |

One of the well-known characteristics of PREs is pairing-sensitive silencing (PSS), where in homozygous conditions, a PRE transgene results in stronger repression as compared to the heterozygous lines (Kassis 1994; Kassis 2002; Kassis AND BROWN 2013). In our assay, >30% of the *ey-*PRE/*ey*-dPRE lines, showed a strong PSS effect in homozygous condition, which is lost in flipped-out lines (Figure 5b, Table 1, and Supplementary Table 4).

In the next step, to ascertain whether the repressor activity of *ey*-PRE/*ey*-dPRE is PcG protein-dependent, we brought a few representative *ey-*PREs lines into mutations background of different PcG proteins by crossing homozygous *ey*-PRE males to the females carrying PcG mutations (Table 2). We compared the de-repression of *mini-white* scored as an increase in the eye pigment level in the progenies from these crosses carrying *ey-*PRE with and without PcG mutation. We found that mutation in *Psc* gene (*Psc^1^* allele) caused a notable de-repression of *mini-*

*white* in five of *ey*-dPRE lines tested. Mutation alleles of *Pleiohomeotic (Pho^1^)* and *Polycomb* (*Pc^1^)* in heterozygous condition exhibited very mild de-repression of *mini-white*. However, a double PcG mutation of *Psc^1^* and *Pc^1^* causes cumulative de-repression effect (Figure 5c, Table 2). Our PRE prediction and ChIP-seq data analysis suggested binding of GAF protein to ey- PRE (Figure 4c). Therefore, we tested GAF mutation (*Trl^R85^*) as well and interestingly loss of GAF caused a notable de-repression (Figure 5c-d and Table 2). These observations suggest that while the function of *ey-*PRE is dependent on PcG genes as in the case of typical PREs, it is possible that factors other than PcG proteins, like GAF, are also involved in its repressive functions. While it was shown for the first time by Hagstrom *et al*. (Hagstrom *et al*. 1997) that GAF mutations effect PRE activity, that GAF is a component of PREs is well documented now (Mishra *et al*. 2001; Mishra *et al*. 2003).

### ME interacts with EB and ey-PRE

As the two CBEs, ME and EB sharply demarcate *ey* locus, we wanted to investigate whether these regions interact in a long-range to mark independent chromatin domain of *ey* regulation. We performed 3C in the 0-16 h embryo using EcoRI restriction enzyme. Using all forward primers to PCR amplify the ligated hybrids (3C DNA) we calculated the relative interaction frequency of ME with several regions at *ey* locus by gel quantification method as described in Naumova et al.(Naumova *et al*. 2012). To assess the primer efficiency, a control template was used (see in Materials and Methods). The fragment comprising ME boundary showed a high interaction frequency with *ey*-PRE and a lesser but still significant interaction frequency with EB and an internal region of *ey* (Figure 6a, 6b iv-vi). However, we did not detect any such interaction of ME with immediate upstream *myo* gene, or further downstream *bt* gene. We also verified the ME and EB anchored interactions in another 3C-qPCR experiment using the restriction enzyme DpnII. This had an additional benefit as DpnII generates smaller fragments containing only a part of ME and EB (Figure 6b iv). The relative interaction frequencies of DpnII fragment containing ME and EB to others were calculated by qPCR as described in Hagege et al. (Hagege *et al*. 2007) using all reverse primer with comparable primer efficiency. We obtained similar results as observed with EcoRI digestion, that ME fragment interacts with *ey*-PRE and EB fragments and it does not interact with upstream region of *myo* or downstream region of *bt* genes. Additionally, EB fragment mainly interacts with ME fragment (Figure 6b v). All the interactions queried are marked with grey looping lines and the ones that tested positive are marked with red looping lines.

**Figure 6.**
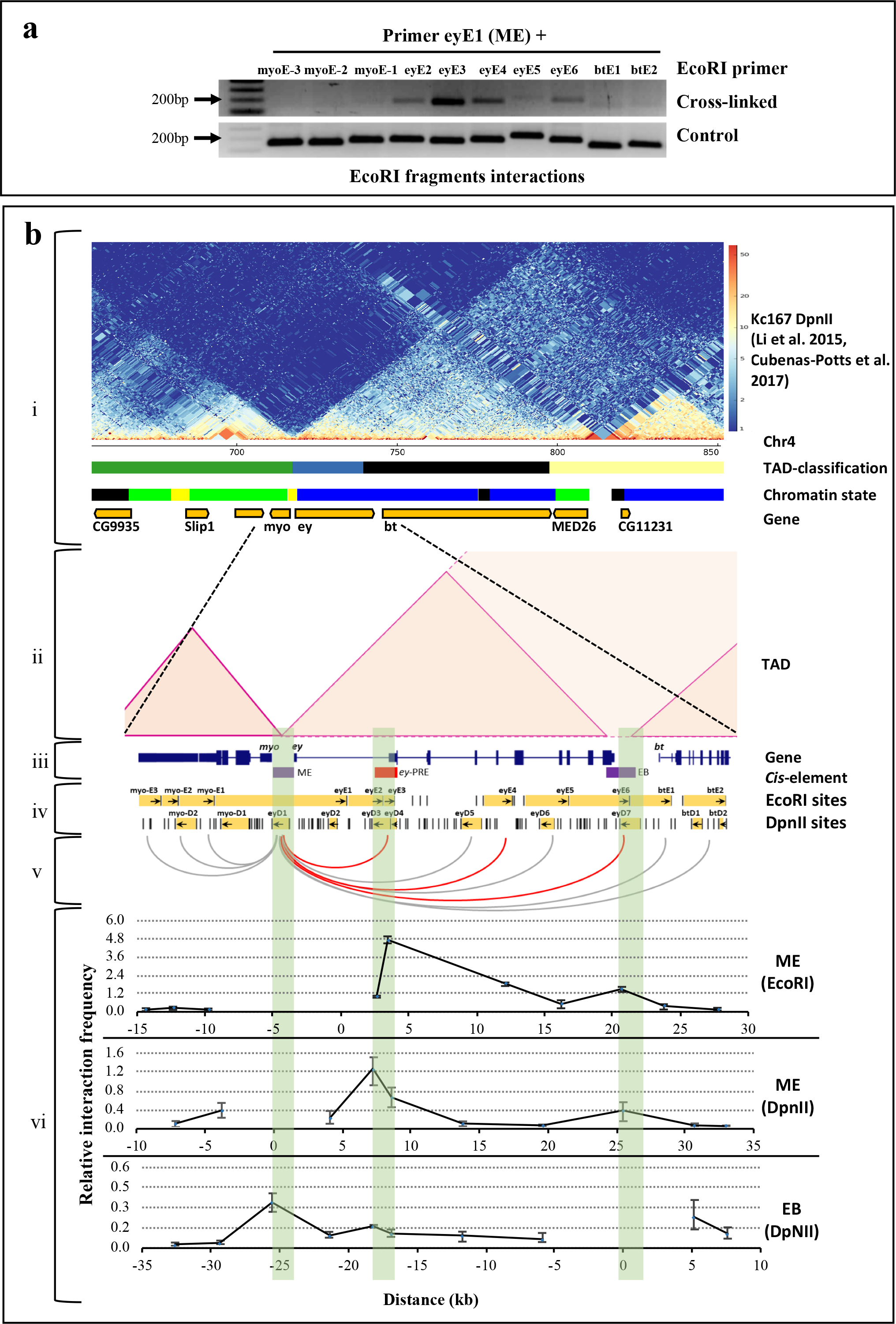
ME, EB, and *ey*-PRE interact in long-range to regulate *ey* locus. **(a)** Interaction of ME with various regions at *ey* locus. 3C was performed in 0-16 h embryo using EcoRI restriction digestion followed by ligation. Ligated hybrids were PCR amplified (cross linked) using all forward primer pairs and confirmed by sequencing. A control template was generated by EcoRI restriction digestion and ligation of equimolar mix of PCR products of all regions flanking to the restriction sites. Primer efficiency of all primers were observed by PCR using the control template (control). Interaction frequencies mean and standard deviations (b-vii, ME- EcoRI) were calculated by gel quantification of three replicates, as described (Naumova *et al*. 2012). EcoRI fragment that contains ME (eyE1) interacts with *ey*- PRE (eyE3) and with an internal region (eyE4) and EB (eyE6) of *ey* (red lines in b-v). However, such interactions are not observed with upstream regions towards *myo* (myoE-3, myoE-2 and myoE-1), other internal regions of *ey* (eyE2 and eyE5) and in farther downstream regions of *bt* (btE1 and btE2) (grey lines in b-v). **(b)** Comparative analysis of chromatin interaction at *ey* locus. (i & ii) A merge Hi-C heatmap of chromatin interactions in ∼250 Kb region of the fourth chromosome along with TAD classification, chromatin states and genes in Kc167 cells. A merged Hi-C data from Li et al. & Cubenas et al. (Li *et al*. 2015; Cubenas-potts *et al*. 2017) was visualized using pyGenomeTracks (Lopez-delisle *et al*. 2021) and chromatin loops were inferred from the same (shaded triangles). The TAD classification track contains the four classifications from ref (Ramirez *et al*. 2018) : Active TAD, yellow; Inactive TAD, black; Inactive TAD contains blue-PcGand green-HP1. The chromatin state track shows five chromatin types from ref (Filion *et al*. 2010) that includes: active chromatin-red and yellow; inactive chromatin-black; PcG-blue; HP1-green. The two genes *ey* and *bt* contain PcG mediated inactive chromatin feature that demarcates a PcG TAD. However, *ey* and *bt* genes appear to be in separate sub-domain within the TAD. (iii) Genes and regulatory region of *ey*. (iv) EcoRI/DpnII sites and uni- directional primers used for 3C. (v) Regions interacting to ME inferred from 3C are marked with red looping lines and non-interacting regions are marked with grey looping lines. (vi) Relative interaction frequencies of EcoRI fragment containing ME to other EcoRI fragments (ME-EcoRI). Relative interaction frequencies of DpnII fragment containing ME (ME-DpnII) and EB (EB-DpnII) with other DpnII fragments . At all the data point’s, mean and standard deviation were generated from 3 experiments. The significant differences to adjacent fragments were assessed by one-way ANOVA using Fisher LSD test, α=0.05. Together 3C and Hi-C data suggest ME and EB interacts in long range to demarcate the *ey* domain.

To understand the *ey* domain organization even better and to validate our 3C results, we explored these interactions in previously published Hi-C data from the whole embryo and a late embryo-derived Kc167 cell, using an online web tool juicebox.js (Sexton *et al*. 2012; Li *et al*. 2015; Cubenas-potts *et al*. 2017; Eagen *et al*. 2017; Robinson *et al*. 2018). A visual inspection of the fourth chromosome Hi-C heatmap from the embryo (Sexton *et al*. 2012) revealed that *ey* and *bt* genes exist in the same TAD, while *myo*, separated by ME boundary is present in a distinct separate domain upstream to *ey* (Supplementary Figure 3). We next looked into the details of chromatin looping within the *ey-*TAD, in high-resolution Hi-C data for Kc167 cells (Li *et al*. 2015; Cubenas-potts *et al*. 2017; Eagen *et al*. 2017). In these data *ey* and *bt* reside in an inactive TAD (*ey*-TAD). Analysis of merged Hi-C data from Li et al. and Cubenas et al. generated in a later TAD classification study (Ramirez *et al*. 2018) suggested that *ey* and *bt* reside in an independent sub-domain within the same TAD (*ey*-TAD). The *ey* lies in a PcG repressed domain while *bt* lies in an inactive domain. However, *myo* and *MED26* are part of separate domains (Figure 6bi and Supplementary Figure 3). The ME boundary appears to be in contact with EB boundary (Figure 6b iv-vi, Supplementary Figure 4). We then compared the expression profile of these genes along with the chromatin states in Kc167 cells to further understand the functional relevance of domain organization at *ey* locus (Filion *et al*. 2010; Cherbas *et al*. 2011). We used the data from a previous study which characterizes the chromatin based on binding of distinct protein combinations and histone modifications (Filion *et al*. 2010). A large region of fourth chromosome has an atypical heterochromatin which is mainly HP1 bound but still permissive to gene transcription. These regions are represented as green chromatin in the chromatin state study. However, we find that the *ey* and *bt* reside in blue chromatin region, a PcG domain, that is distinct from the flanking green chromatin regions (Figure 6bi). In concordance, *ey* and *bt* are expressed at a very low level suggesting their co-regulation in a transcriptionally repressed domain. While, *myo* and *MED26* falling in green chromatin region show a higher level of expression (Supplementary Figure 3b).

In addition, we also compared the interactions of ME and EB in antennal disc and eye disc from third-instar larvae using Hi-C data from Viets et al. (Viets *et al*. 2019). In third instar larvae, the expression of *ey* is restricted to anterior of morphogenic furrow in eye disc while it is not expressed in antennal disc altogether. In both tissues, *ey*-TAD remains the same as in Kc167 cells. A similar level of interaction between ME and EB was observed in both the tissues, however, interaction of ME to *ey*-PRE was more prominent in eye disc (Supplementary Figure 4b). These observations together suggest a universal interaction of ME with EB in the embryo and Kc167 cells and this reflects the role of the two CBEs in defining a distinct chromatin domain. The interaction of ME boundary with *ey-*PRE is more functionally determined.

### ME, EB and ey-PRE associate differently with nuclear matrix

Nuclear Matrix (NuMat) has been proposed to provide a structural framework for targeted tethering of chromatin domain in sub-nuclear space (Mishra AND KARCH 1999). DNA that associates with NuMat (MARs) are proposed to be the anchoring sequence. CBEs have been shown to interact with each other and along with boundary proteins are shown to associate with NuMat (Blanton *et al*. 2003; Byrd AND CORCES 2003; Pathak *et al*. 2007). As we observed interaction amongst ME, EB, and *ey*-PRE, we further investigated whether these sequences associate with NuMat using an *in vivo* MAR assay (Mirkovitch *et al*. 1984) (Supplementary Figure 5). In the assay, equal amount of PCR amplified test regions was Southern hybridized with radio-labelled NuMat DNA isolated from *Drosophila* embryos. NuMat association is determined by comparative quantification of hybridization signal of test regions with known MAR in Histone gene (HIS-MAR – positive control) and a non-MAR region in exonic sequence of the BEAF-32 gene (BEAF-CDS – negative control) (Mirkovitch *et al*. 1984; Pathak *et al*. 2007).

We found that ME and EB both show a comparable level of association with NuMat. Interestingly, the *ey*-dPRE and the EB-u (3’-UTR region) of *ey* gene also show a strong association with NuMat. However, the INT7 region does not show significant association with NuMat (Figure 7a-c). Our NuMat binding assay using embryo had limitations as during embryonic development, *ey* is expressed in a limited number of cells, i.e., eye-antennal disc primordium and embryonic brain cells (Hauck *et al*. 1999). Reasoning that we might be losing important information as *ey* does not express in over-whelming majority of embryonic cells, we also tested the NuMat association of the locus in a different tissue with significant *ey* transcriptional activity, i.e., brain-eye-antennal imaginal discs of third instar larvae. In third instar larvae, *ey* mainly expresses in CNS and anterior to the morphogenetic furrow in the eye disc. As an other extreme, we also tested NuMat association of *ey* locus in S2 cells where the gene is not transcribed (Czerny *et al*. 1999; Hauck *et al*. 1999; Adachi *et al*. 2003; Chintapalli *et al*. 2007). The NuMat associations of ME, *ey-* PRE, and EB varied in the embryo, brain-eye-antennal disc and S2 cells with the association being stronger when the gene is expressed. Interestingly, *ey*-PRE was found to associate with NuMat in all the tissues examined, irrespective of *ey* gene transcription status. ME and EB boundaries were found to associate with NuMat in embryos and brain-eye disc, while the association was completely absent in S2 cells. EB-u association to NuMat is stronger than EB-i, indicating that while EB-i harbours boundary activity, EB-u is responsible for tethering of the locus to nuclear architecture (Figure 7).

**Figure 7.**
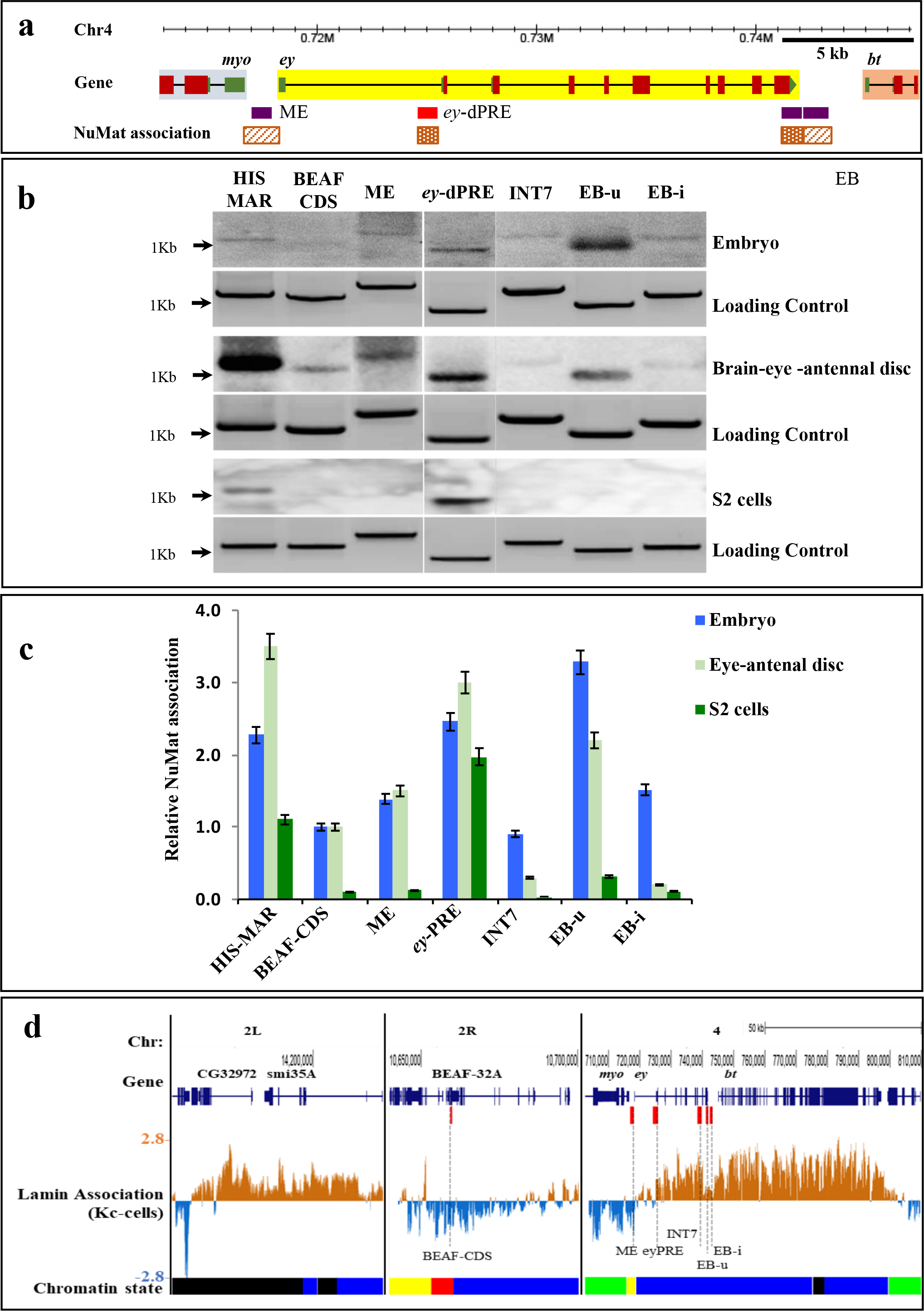
Different regions of *ey* associate with NuMat. **(a)** A map of *ey* locus along with NuMat association of *cis-*elements ME, *ey-*dPRE, EB-u, and EB-i. **(b)** NuMat association of *ey cis-*elements along with known MAR (histone MAR region – HIS MAR) as a +ve and BEAF coding sequence (BEAF CDS) as a -ve control. The PCR products of test and controls DNA (∼400ng) were electrophoresed on 1% agarose gel, transferred to the NY+ membrane. Southern hybridization was carried out with radiolabeled NuMat DNA from the embryos (0-16 h), brain-eye-antennal disc (3rd instar larvae), and S2 cells. Initial gel image of electrophoresed PCR products before the southern hybridization was used as loading control. After processing of the gel image, lanes have been spliced and arranged in order for clarity. **(c)** Relative NuMat association of all test regions were calculated by quantitation of the signal band from blot/gel using ImageJ. The signal intensities were normalized over the loading controls and then compared with the negative control, BEAF-CDS. The BEAF CDS shows some basal level of NuMat association that has been taken into consideration for normalization. Signal intensity of the negative control was considered at value one in embryos. The mean values and standard deviation were calculated using three independent experiments in embryos, eye-antennal disc and two independent experiments in S2 cells. *Ey-*dPRE and EB-u show a strong NuMat association (comparable to HIS MAR) in the embryo and brain-eye-antennal disc. ME and EB-i show weak interaction to the NuMat in the embryos while in brain-eye-antennal disc EB-i does not appear to interact with the NuMat . In S2 cells, only e*y-*dPRE associates with the NuMat whereas in all experiment INT7 does not interact with NuMat. **(d)** Lamina association in *ey*-TAD region. Lamin binding at known Lam target region-CG32972, BEAF-CDS (negative control), and *ey*-TAD regions using Lam-DamID data from ref (Filion *et al*. 2010). The chromatin state track shows five chromatin types that includes: active chromatin-red and yellow; inactive chromatin-black; PcG-blue; HP1-green. The genes *ey* and *bt* contain PcG mediated inactive chromatin and associate with the nuclear lamina.

Nuclear lamina is a part of the NuMat and mostly repressive in nature. As we observed that *ey* and *bt* are present in a sub-domain of the *ey*-TAD with repressed chromatin features (a PcG bound region) we queried whether they reside in a lamina associated domain. We used *Drosophila* genome-wide binding data for lamin-B in Kc167 cells generated by DamID (Filion *et al*. 2010). Interestingly, we found both the genes were targeted to lamina with intensities comparable to a known lamina targeted gene, i.e. CG32972 (Pickersgill *et al*. 2006). The lamina association towards *ey* region was comparatively lower than that of *bt* region. Interestingly, the lamina association of ME, EB and INT7 was lower compared to *ey*-PRE. This observation agrees well with our data where we find that in S2 cells, these regions are not bound to NuMat, while *ey*-PRE does bind to NuMat. Altogether these observations suggest that association of *ey* and *bt* to lamina may be responsible for organizing the *ey*-TAD in a PcG meditated repressed domain in cells where the genes are not expressed. Further, the *ey cis-*regulatory elements associate differentially to NuMat depending on the transcriptional activity of the locus. While these observations provide an initial clue to the 3D organization of the locus, detailed studies are needed to fully understand the role of MAR sequences in dynamics of the region.

## Discussion

The *ey* gene crucial for eye development in *Drosophila*, is differentially expressed in comparison to its neighbouring genes, *myo,* and *bt*. Transcriptional regulation of *ey* is a challenge, as the flanking genes and their regulatory sequences are very close to it. Specifically, the *myo* is 1.6 Kb upstream and *bt* is 3.2 Kb downstream. (Desplan 1997; Halder *et al*. 1998; Pichaud AND DESPLAN 2001). Earlier, we have identified ME boundary in the *myo* and *ey* intergenic region (Sultana *et al*. 2011). Based on the distinct expression pattern of the three genes, we propose that additional *cis*-regulatory elements like CBEs and PREs might be present at this locus. In order to identify such elements and to understand their potential role in *ey* regulation, we used bioinformatics prediction tools and clusters of binding sites of known CBE and PRE binding proteins from genome-wide localization studies. We further co-mapped the region to DHSs that are often indicative of the presence of regulatory elements. Taking clues from *in silico* prediction, we have identified and functionally characterized a novel CBE and a PRE associated with *ey* locus. We show that ∼1.2 Kb intergenic region between *ey* and *bt*, EB, functions as boundary and ∼1.6 Kb region, upstream to *ey* promoter, *ey*-PRE, functions as PcG dependent repressor.

Our *in silico* analysis estimates three putative CBEs in *ey* locus, i.e., ME, INT7 and EB. Of these, ME boundary has already been characterized earlier. In the present work, we report that only EB displays a resolute boundary activity during embryonic development as well as a mild boundary activity in the adult stages. Whereas, INT7 shows a scant boundary activity only during early embryogenesis suggesting that boundary activity of these elements could be developmentally regulated. A few recent studies have suggested a developmental and tissue-specific regulation of CBEs mediated by spatially and temporally controlled expression of boundary proteins (Aoki *et al*. 2008; Lin *et al*. 2011; Aoki *et al*. 2012; Matzat *et al*. 2012; Bonchuk *et al*. 2015; Wolle *et al*. 2015; Gambetta AND FURLONG 2018). For example, expression of boundary factors, Elba and a late boundary complex (LBC), mediate boundary functions of Fab7 element in early embryogenesis and adults, respectively (Aoki *et al*. 2008; Aoki *et al*. 2012; Wolle *et al*. 2015). In another example, a tissue-specific protein Shep has been shown to modulate *gypsy* boundary activity and nuclear localization, particularly in CNS (Matzat *et al*. 2012). Our study of *ey* locus presents a similar scenario, where the boundaries are spatially and temporally regulated depending on the transcriptional activity of the locus. It would be further interesting to see if ME, EB and INT7 functions are developmentally regulated or restricted to specific cell-types.

Previous studies have shown that interaction of BEAF-32 with CP190 mediates long-range chromosomal contact, and more than 70% of BEAF-32/CP190 bindings demarcate domain boundaries in the *Drosophila* genome (Vogelmann *et al*. 2014; Cubenas-potts *et al*. 2017; Wang *et al*. 2018). The EB element also harbours *in vivo* binding sites of BEAF-32, CP190, and CTCF co-mapping with a prominent DHS. This overall architecture of the EB element shows a remarkable similarity to the earlier identified ME (Sultana *et al*. 2011). These evidences suggest that the same molecular players, particularly BEAF-32 and CP190, might be contributing to the functional linking of the two CBEs to define the loop domain of *ey*. In our 3C experiment, we confirm the interaction between ME and EB suggesting that the two CBEs, flanking the *ey* locus, define functional autonomy of the chromatin loop domain. In support to our finding, we observe the chromatin loop of ME with EB in Hi-C data from previous studies in Kc167 cells which place *ey* in an independent domain within a larger TAD (Li *et al*. 2015; Cubenas-potts *et al*. 2017; Eagen *et al*. 2017). A similar and comparable interaction between ME and EB is also seen in tissues with differential *ey* expression. (Ref) These interactions provide necessary insulation to *ey* locus from the neighbouring regulatory environments.

We report here the identification and functional validation of a new CMM or PRE, *ey*- PRE, within the *ey* domain. Besides conferring a repressive feature in the transgene-based assay, the element also exhibits PSS. The *ey-*PRE encompasses a core DHS region spanning ∼830 bp (*ey-*dPRE) that coincides with sharp binding peaks of PcG proteins of PRC1 complex, namely, Pc, Psc, dRing. However, in the transgenic assay, only Psc mutation shows a prominent de-repressive effect. Mutations in other PcG proteins show only a moderate impact on *ey-*PRE activity. One reason for this variation could be that these mutations were tested in heterozygous condition, as homozygous mutants are lethal. In case of double PcG mutants however we could see a strong de-repressive effect. In summary, the results suggest that *ey*-PRE maintains the expression state of the *ey* locus and requires the activity of a subset of PcG proteins for the purpose. The properties of *ey-*PRE is similar to the earlier reported PREs (Kassis 2002; Kassis AND BROWN 2013). Interestingly, a non-PcG protein, GAF, was found to attenuate the function of *ey*-PRE. We have earlier reported that ME boundary interacts with GAF protein (Sultana *et al*. 2011). This tempts us to speculate that the observed 3C interaction of ME and *ey-*PRE could be mediated by GAF protein. Recent Hi-C studies also support the interaction of PREs with multiple regions within a TAD and active role of GAF in such connections (Ogiyama *et al*. 2018). The *ey-*PRE also has a predicted promoter property, which suggests it might have a dual role of promoter as well as CMM. However, the role of GAF remains unexplored and it will be interesting to see if GAF is crucial to these diverse functions of same DNA segment.

Studies have shown that NuMat provides anchoring sites for the compartmentalization of higher-order chromatin organization and also acts as a platform for functional activities inside the nucleus. Work from our lab and elsewhere have shown that boundary elements function in the context of NuMat (Byrd AND CORCES 2003; Pathak *et al*. 2007). In this study, we show that multiple sites of the *ey* locus, including ME, *ey*-PRE, and EB, associate with NuMat in transcription status dependent manner. As BEAF-32 is a bona fide NuMat associated protein in flies, and ME and EB, both carry BEAF-32 binding sites, the anchoring of these CBEs to the nuclear architecture is probably mediated by the protein (Pathak *et al*. 2007). The prominent association of ey-PRE to NuMat in all cell types, unlike other elements, suggests the role of the memory element in all functional states. Taken together, the dynamic nature of NuMat association of different regulatory elements of the locus supports the hypothesis that *ey* locus is present in different compartments depending on the expression state of the gene and that NuMat might be providing the structural basis for this compartmentalization.

Based on these findings, we propose a model for *ey* gene expression where *ey*-PRE functions as both PRE/TRE to maintain the active or repressed state of *ey* in different cell/tissues initially established by early developmental cues. The CBEs, ME and EB, demarcate the *ey* locus, to create an independent domain of differential regulation, insulated from the regulatory elements of neighbouring genes *myo* and *bt.* The *ey*-PRE, along with the other unidentified regulatory elements of *ey* locus, which are bound to PcG or trxG proteins, probably takes the *ey* domain either to repressive (*Polycomb* body) or active (transcription factory) compartment, respectively (Figure 8).

**Figure 8.**
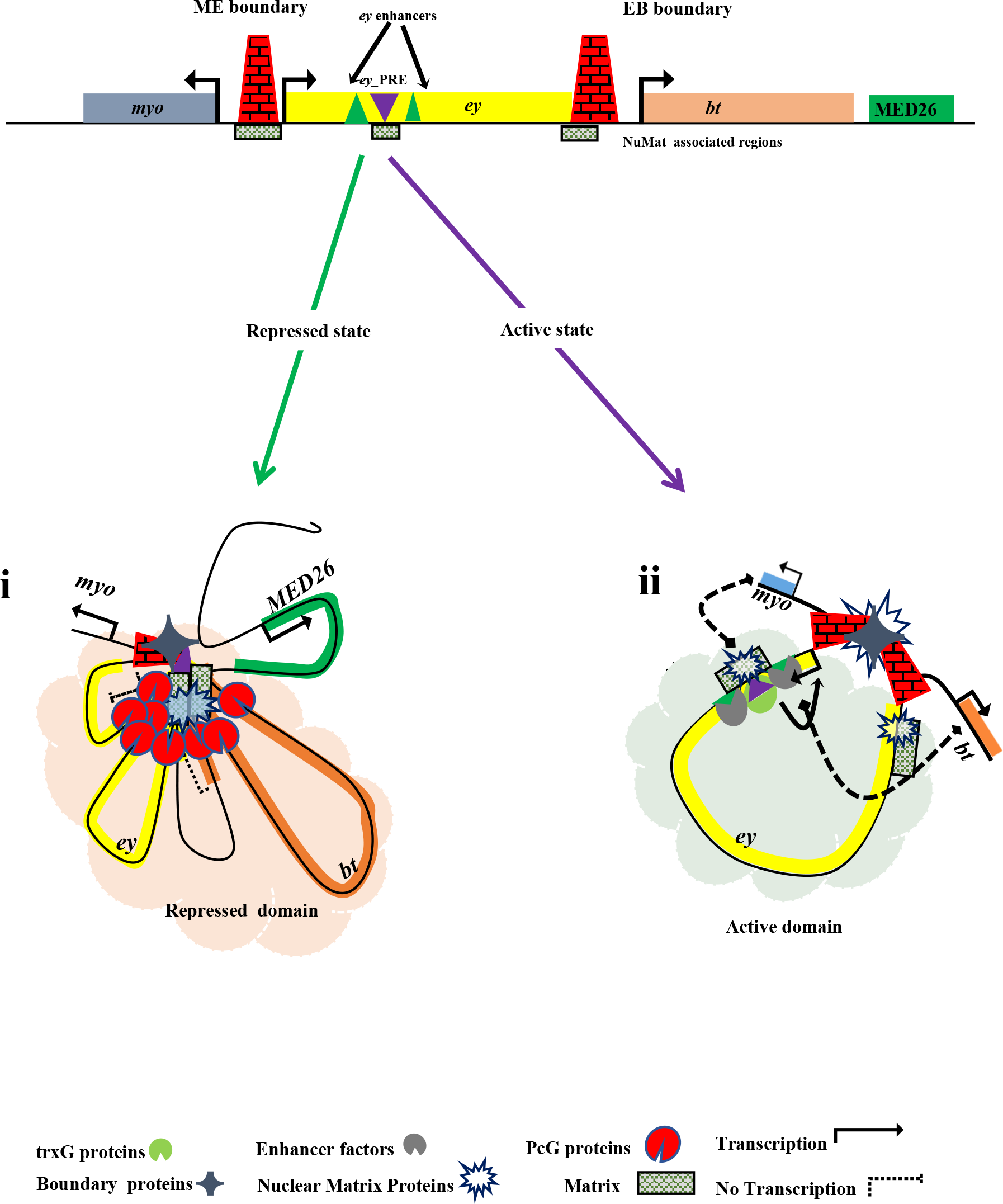
A Model for regulation of *ey* locus. In the absence of activation signal by initial regulatory proteins or upon receiving a repressive signal during development, PcG proteins bind to *ey-*PRE to bring and maintain the transcriptional repression of *ey*. As *ey* and *bt* are transcriptionally inactive and present in a common TAD and associate to nuclear lamina in Kc167 cells, it is likely that *ey* and *bt* occupy the same repressed nuclear space. Further, the tethering of *ey* and *bt* regions to NuMat and the chromatin interactions of ME with *ey-*PRE, EB, and downstream regions of the *bt* collectively determine the higher-order chromatin organization at the repressed domain (**i**). Upon activation signal during development, the *ey* gene moves out into an active compartment of the nucleus likely with the help of *ey*-PRE which might now function as a TRE and associate with NuMat for activity (**ii**). Moreover, *ey* promoter that coincides with *ey*-PRE becomes a target of activators, i.e., trxG proteins and enhancer proteins to activate *ey*. At the same time, ME and EB boundaries demarcate the independent domain of *ey* that restrict cross-regulatory influences with neighbouring regulatory elements of genes *myo* and *bt*.

### Conclusions

Our work defines the *Drosophila ey* domain demarcated by two CBEs. Fine dissection of the locus revealed a CMM involved in maintaining the expression status of the locus. These regulatory elements interact among themselves and with the nuclear architecture depending on the transcriptional status of *ey* gene. Further studies using super-resolution imaging of the locus in a cell-type-specific manner and identification of molecular players involved in the long-range interactions of these regulatory elements will provide a deeper insight into the regulation of the locus. At present, our study provides clues to the hierarchical organization of the chromatin that form the basis of spatiotemporally regulated gene expression in euchromatin.

## Methods

### CBEs and PRE prediction in ey locus

A ∼ 30 Kb *ey* region was used in cdBEST analysis to identify new CBEs at *ey* locus. Additionally, for prediction of the CBEs, *in*-*vivo* binding of known boundary proteins BEAF32, CP190, CTCF, and GAF was analyzed using whole embryo (0-12 h) ChIP-chip data from a previous by study Negre et al., 2010 (Negre *et al*. 2010). We co-mapped the DNaseI hypersensitive sites(DHS) in the region using stage-9 (∼ 6 h) embryo DHS data available at UCSC browser (https://genome.ucsc.edu/) (Thomas *et al*. 2011). Two potential CBEs, a ∼ 1.3 Kb intronic region in a seventh intron (INT7) and a ∼ 2.2 Kb intergenic region between *ey* and *bt* (EB), were identified that also co-map with DHS (Figure 1).

A Perl based program, PRE mapper, was used to predict putative PRE (*ey-*PRE) in the *ey* locus (Srinivasan AND MISHRA 2020). PRE mapper analyzes motifs of known PRE binding proteins (Pho, GAF, Dsp1, Sp1, Zeste, Grh, Adf1, and Cg), their binding patterns, and motif clustering to define a PRE. Presence of DHS at *ey*-PRE was reviewed to correlate its regulatory feature. Additionally, a comprehensive analysis of various chromatin features in the *ey* region was carried out using 14-16-h whole embryo ChIP-seq data available at modENCODE by Gary Karpen. modENCODE id of the data used are H3K27Ac (4120), H3K27Me3 (3955), Pc (3957), dRING (5071), Psc (3960), BEAF-32 (3954), CP190 (3959), CTCF (5069), and GAF (4149) (Mod *et al*. 2010).

### Cloning of predicted CBEs and PRE

All the test fragments were PCR amplified from *CantonS*(*CS*) fly genomic DNA by high fidelity *Pfu* polymerase with specific primers (Supplementary Figure 1, Supplementary Table 1 and 6). All CBE test fragments (INT7, EB, EB-u and EB-i) were inserted first at MCS in pLML and then sub-cloned along with *loxP* at XhoI sites in the enhancer-blocker assay vectors, pRW+ and pCfhL, (Sultana *et al*. 2011). Test PREs (*ey*-PRE and *ey*-dPRE) were cloned initially at SmaI blunt site in pLML, then sub-cloned along with *loxP* at XhoI in pCasPerX (Vasanthi *et al*. 2013).

### Fly culture, transgenic flies, and genetic crosses

All fly strains of *Drosophila melanogaster* were cultured in standard cornmeal medium at 25 ^0^C. Transgenic flies were generated by micro-injection of an assay vector DNA and a transposase expressing plasmid *p* Δ *2-3* together into pre-blastoderm of *w^1118^* as described in Siegal et al. (Siegal AND HARTL 2000). Adult flies from injected embryos were backcrossed to *w^1118^,* and the transgenes were identified by the presence of eye pigment in the adult progenies. Transgenic flies were next crossed with the double balancer (*Pin/CyO; TM2/TM6*) to know chromosomes of P-element integration and making homozygous stock. We established several lines for each predicted CBE and PRE (Supplementary Table 2-4). The test fragment was flipped out from initial transgene, to compare the effect of test fragment and rule out any position effect, by crossing the flies with *Cre-recombinase* expressing flies and later was confirmed by PCR using specific primers. The homozygous *ey*-dPRE males were crossed to virgin females of different PcG proteins mutant (*Pc^1^, Psc^1^, Pho^1^*) to investigate the effect on repressor function of *ey*-dPRE (listed in Supplementary Table 7). The effect of PcG mutations effect were compared in progenies with or without mutations in heterozygous condition.

### Visualization and quantitative measurement of eye pigment

The level of pigment in the eye of similar aged (5-day) adults were visually determined to compare the level of *mini-white* expression in the enhancer-blocker and PRE assay transgenic flies. In transgenic flies, pigmentation range varied from bright/dark red in wild-type (*CS*) toward red, dark orange/light red, yellow, and light-yellow, to white in the absence of expression (Supplementary Table 2-4). We also quantitated the red pigment value in eye for some transgenic flies, to compare the strength and effect of boundary and PRE after flipping-out (Sultana *et al*. 2011). Eye pigments from a minimum of ten adult heads were extracted in a 1:1 mixture of chloroform/0.1% ammonium hydroxide (200 µl) then centrifuged the mix for 2 min at 13000 rpm. Finally, 100 µl of the supernatant was taken for quantification of red pigment by spectrophotometric absorbance at 485 nm. For each genotype, a mean relative pigmentation value and standard deviation were calculated from a minimum of three independent experiments compared to the pigment value of *w^1118^*.

### **β**-galactosidase staining and quantification

For enhancer blocker assay in embryo, lacZ staining was performed to determine the beta-galactosidase (lacZ) activity (Bellen *et al*. 1989; Sultana *et al*. 2011). In brief, 0-12-h old embryos were collected and dechorionated in 50% bleach (sodium hypochlorite). The dechorionated embryos were washed and fixed for 20 min in heptane (saturated with 25% glutaraldehyde). The fixed embryos were again washed thoroughly with 1x PBST (0.3% Triton X-100), incubated initially for 10 min at 37 ^0^C in lacZ staining solution (3 mM K_3_[Fe(CN)_6_], 3 mM K_4_[Fe(CN)_6_], 7.2 mM Na_2_HPO_4_, 2.8 mM NaH_2_PO_4_, 1 mM MgCl_2_, 150 mM NaCl), and finally stained for 4-6 h in lacZ staining solution with 0.2% X-gal (Sigma). The stained embryos were imaged in Leica stereomicroscope. We performed lacZ staining of multiple transgenic lines along with positive and negative controls in a common grid, ensuring reliable comparison of staining between different genotypes.

To estimate and compare the enhancer blocker activity of the test fragments in the embryos, we quantitated lacZ stained embryos using ImageJ-image processing and analysis software provided by the National Institutes of Health, USA (Hartig 2013). To calculate relative lacZ staining and standard deviation and plot it as a bar graph for comparison, we used mean pixels value of a fixed area size of different regions from multiple embryos (3-5) of each genotype. The embryos represented overall staining pattern.

### 3C-PCR and qPCR

Wild type *CS* embryos (0-16h old) were processed for 3C, using EcoRI and DpnII separately, as described earlier (Lanzuolo *et al*. 2007; Comet *et al*. 2011) with few modifications. To begin with, 1 g embryos were dechorionated in 50% fresh bleach for 2-3 min and washed thoroughly with PBST (PBS 1x + 0.01% Triton X 100). The dechorionated embryos were fixed for 15-20 min at room temperature (RT) with 2% formaldehyde (Sigma) in 5 ml fixing solution (50 mM HEPES-pH-7.6, 100 mM NaCl, 0.1 mM Na-EDTA-pH-8, 0.5 mM Na-EGTA-pH-8) added with 15 ml heptane. To stop fixation, formaldehyde was quenched by 0.125 M glycine for 5 min at RT. The fixed embryos were resuspended in 2.5 ml of ice-cold lysis buffer (10 mM Tris-pH-8, 10 mM NaCl, 0.2% NP40 with Roche protease inhibitor cocktail freshly added) and homogenized in Dounce homogenizer (10-15 strokes) to create nuclear suspension. Cellular debris was removed from the nuclear suspension by filtration through a double layer of Mira cloth. Nuclei were pelleted at 5000 rpm for 5 min, then washed once and resuspended in 500μl 1.2x EcoRI or DpnII restriction enzyme buffer from New England Biolabs (NEB). An aliquot of 50μl of nuclear solution (∼100mg embryos) was diluted in 362μ 1.2x restriction enzyme buffer and further used for restriction digestion. Sample was sequentially treated with 0.3% SDS for 1 h at 37^0^C and then 2% triton X-100 for 1 h at 37^0^C with continuous mixing at 1000 rpm. Subsequently, restriction digestion was carried out using 400 units of EcoRI (NEB) or 1500 units of DpnII (NEB) for 2 h at 37^0^C and continuous mixing at 1000 rpm. Restriction enzyme was heat inactivated with SDS added to a final concentration of.5 % and heated at 65^0^C, mixed at 100 rpm for 20 min. At this step, 100 μl of digested sample was taken aside for digestion efficiency calculation. The remaining sample was diluted in 10 ml 1x T4 DNA ligase buffer (NEB) containing 1% Triton X- 100 and incubated at 37^0^C and 750 rpm for 1 h. Ligation was performed for 4 h at 22^0^C, mixing at 750 rpm with 40000 unit of T4 DNA ligase (NEB). The ligated sample (3C DNA) and control samples (fixed DNA and digested DNA) were sequentially treated with RNaseA (100 μg/ml) at 37^0^C for 1 h and with Proteinase-K (500 μg/ml) at 55^0^C for 1 h followed by overnight de-crosslinked overnight at 65^0^C. The 3C DNA and control DNA were extracted by phenol: chloroform: isoamyl alcohol (25:24:1) and ethanol precipitation method. Concentration of extracted DNA samples were quantified using Qubit 2.0 fluorometer (Thermo Fisher Scientific) and adjusted to 50 μng/ l.

For each 3C experiment efficiency of digestion and ligation were visually inspected on agarose gel by comparing equal amount of fixed DNA (control), digested DNA and ligated DNA (3C DNA). Additionally, digestion efficiencies of several restriction sites at *ey* locus were also calculated using PCR (for EcoRI) and qPCR (for DpnII) as described (Hagege *et al*. 2007; Naumova *et al*. 2012). The 3C DNA from experiments with 80% and above digestion efficiency were used for interaction frequency analysis. Since we did not have BAC clone available for *ey* region, we generated control DNA templates for primer efficiency estimation. We used forward-reverse primers pair to PCR amplify the ey genomic regions at several EcoRI/DpnII restriction sites. PCR amplified regions were mixed in an equimolar ratio, restricted digested, and ligated. Unidirectional primer pairs were design for interaction frequency analysis. Quality and efficiency of all the primers were checked with control DNA template generated. Primers with very low efficiency or generating more than one amplicon were not used further with 3C DNA analysis.

For EcoRI 3C, interaction of ME containing EcoRI fragments to different regions were checked by PCR on 100 ng 3C DNA using all forward primers paired to ME forward primer-eyE1 (Supplementary Table 6). All PCRs amplifications were carried out with the following parameters: 95^0^C for 3 min, followed by 36 cycles of 95^0^C- 30 s, 56^0^C-10 s, and 72^0^C-8 s, with a final step at 72^0^C-2min. The 3C DNA PCR products were resolved on 1.2% agarose gel, and we confirmed the chimeric sequences by sequencing of gel extracted PCR DNA product. The signal intensity of the 3C PCR product in the gel was quantified using ImageJ. Finally, relative interaction frequency and standard deviations for each forward primer pairs were calculated from three replicates as described in Naumova et al. (Naumova *et al*. 2012). For DpnII 3C, interaction of ME and EB fragments with others were assessed by qPCR on 100 ng 3C DNA using all reverse primers paired with ME reverse primer (eyD1) and EB reverse primer (eyD7), respectively (Supplementary Table 6). All qPCR reactions were set using Power SYBR® Green PCR Master Mix (applied biosystem) using following conditions: 95^0^C for 10 min, followed by 45 cycles of 95^0^C 15 s, 60^0^C for 60 s (acquisition). We calculated the relative interaction frequency as described in Hagege et al.(Hagege *et al*. 2007) normalized over loading control using an EcoRI site primer pair (non DpnII restriction site).

### Nuclear matrix Association Assay

The nuclear matrix (NuMat) association of *ey cis*-elements were tested in a modified *in vivo* MAR assay from the original protocol of Mirkovitch, *et al*. 1984 (Mirkovitch *et al*. 1984; Pathak *et al*. 2014) (Supplementary Figure 3).

First, NuMat DNA was prepared from 0-16 h old embryos, eye-antennal discs (3rd instar larvae), and S2 cells. In brief, nuclei were isolated from 1 g of embryos and ∼ 200 pairs of eye-antennal imaginal discs and 2x10^6^ S2 cells. An aliquot of nuclei was used for the isolation and estimation of total genomic DNA for quality control checks. Isolated nuclei were treated with DNaseI in nuclei isolation buffer (20 mM Tris-pH- 7.4, 20 mM KCl, 70 mM NaCl, 10 mM MgCl_2_, 0.12 5mM spermidine 1mM PMSF, 0.5% Triton-X 100, and 200μg/ml DNaseI) at 4^0^C for 1 h to remove chromatin.

Chromatin depleted nuclei were collected by centrifugation at 3000 x g for 10 minutes. Non-matrix proteins were extracted out from the nuclei with 0.4 M NaCl for 5 min in extraction buffer (10 mM Hepes-pH-7.5, 4 mM EDTA, 0.25 mM spermidine, mM PMSF, 0.5% (v/v) triton X-100) and another 5 min with 2 M NaCl in the extraction buffer to get NuMat. The NuMat pellet was washed twice with wash buffer (1 mM Tris-pH-7.4, 20 mM KCl, 1 mM EDTA, 0.25 mM spermidine, 0.1 mM PMSF) then treated with RNaseA (20 μg/ml) for 30 minutes at 37^0^C and with 100 μg/ml Proteinase-K at 55^0^C for 1 h. Finally, NuMat DNA was obtained by phenol:chloroform:isoamylalcohol (25:24:1) and ethanol precipitation method. NuMat DNA was dissolved in DNase free water and quantified using Nano-drop. An equal amount of NuMat DNA isolated from different tissues was labeled with 32^P^-dATP by the Random Primer Labeling method.

Next, all the test regions were PCR amplified from fly genomic DNA using specific primers listed in Supplementary Table 6. A known histone gene matrix-associated region (HISMAR) was used as a positive control and an exonic sequence of BEAF- 32 gene (BEAF CDS) was used as a random control. The controls were used to compare the relative NuMat associations. An equal amount (∼ 400 ng) of all PCR products were resolved first on a 1.2% TAE agarose gel and then transferred to a positively charged nylon membrane using a capillary transfer method. The membrane was then subjected to a standard Southern hybridization using ^32^P-dATP labeled NuMat DNA as described earlier (Pathak *et al*. 2014). Finally, the hybridization signals were observed in Phosphor Molecular Imager (PMI) from (BIORAD). Relative association to NuMat and standard deviation for each region was calculated by signal intensity from three replicate blots using ImageJ (Hartig 2013).

### Analysis of Hi-C, chromatin states and lamin Dam-ID data from previous studies

To understand *ey* locus organization, we analyzed processed Hi-C matrix data from previous studies (Supplementary Table 5) using two online visualization tools, i.e., Juicebox (Robinson *et al*. 2018) and HiCExplorer/pyGenomeTracks (Wolff *et al*. 2018; Lopez-delisle *et al*. 2021). The processed Hi-C data of embryonic stage from Sexton et al. (Sexton *et al*. 2012) and Kc167 cells from Eagen et al. (Eagen *et al*. 2017) were visualized in Juicebox. We also used a high resolution merged Hi-C data from Kc167 cells from Li et al., and Cubenas et al. (Li *et al*. 2015; Cubenas-potts *et al*. 2017) available online at chorogenome.ie-freiburg.mpg.de (Ramirez *et al*. 2018). The processed merged Hi-C matrix, TAD classification (Ramirez *et al*. 2018) and chromatin states data (Filion *et al*. 2010) were visualized using HiCExplorer/pyGenomeTracks tools available publicly at European Galaxy server (hicexplorer.usegalaxy.eu). Finally, we used Hi-C data from Kc167 cells, antennal disc and eye disc (Viets *et al*. 2019) for comparative analysis of TAD boundaries and relative interaction of ME, *ey*-PRE and EB regions at *ey* locus.

To know lamina association of *ey* domain and its chromatin features, we used chromatin states and lamin Dam-ID data of Kc167 cells from Filion et al. (Filion *et al*. 2010). For comparison, we visualized the data at a known lamina associated region- CG32972 (Marshall *et al*. 1996; Pickersgill *et al*. 2006), BEAF coding region and *ey* locus.

### Statistical analysis

We performed analysis of variance (ANOVA) using Graph-pad prism software to statistically analyze the data. A one-way ANOVA with multiple comparisons test (Sidak’s/Fisher LSD, α=0.05) was applied to compare the means, standard deviations and to detect significant differences between the groups for relative pigment values in adult eyes, lacZ staining in embryos and NuMat associations. In 3C, the mean and standard deviation of relative interaction frequency of each restriction fragment and the significant differences to adjacent fragments were assessed by one-way ANOVA using Fisher LSD test, α=0.05.

## CRediT author contribution statement

**Shreekant Verma:** Investigation, Methodology, Visualization, Writing – original draft.

**Rashmi U Pathak**: Investigation, Writing – review and editing.

**Rakesh K Mishra**: Resources, Visualization, Supervision, Funding acquisition, Writing – review and editing.

## Acknowledgements

We thank Welcome Bender for his discussion and sharing of Pc binding data in *ey* locus (not shown). We thank Jaya Krishnan for helping us in lacZ staining of the CfhL transgenes. We thank Rahul Sureka for helping in eye-antennal disc isolation. We also thank Arumugam Srinivasan for his helpful inputs in PRE predictions and Yaser Payferman for his help in cloning for *ey*-PRE.

## Data availability

The processed data from ChIP-chip, DHS, ChIP-Seq, Hi-C samples and Dam-ID obtained from online sources are summarized in Supplementary Table 5. Fly mutants or transgenic lines and their source are listed in Supplementary Table 7. Transgenes generated in this study are available on request.

## References

Adachi, Y., B. Hauck, J. Clements, H. Kawauchi, M. Kurusu et al., 2003 Conserved cis-regulatory modules mediate complex neural expression patterns of the eyeless gene in the Drosophila brain. Mech Dev 120: 1113–1126.

Ahanger, S. H., A. Srinivasan, D. Vasanthi, Y. S. Shouche and R. K. Mishra, 2013 Conserved boundary elements from the Hox complex of mosquito, Anopheles gambiae. Nucleic Acids Res 41: 804–816.

Americo, J., M. Whiteley, J. L. Brown, M. Fujioka, J. B. Jaynes et al., 2002 A complex array of DNA-binding proteins required for pairing-sensitive silencing by a polycomb group response element from the Drosophila engrailed gene. Genetics 160: 1561–1571.

Aoki, T., A. Sarkeshik, J. Yates and P. Schedl, 2012 Elba, a novel developmentally regulated chromatin boundary factor is a hetero-tripartite DNA binding complex. Elife 1: e00171.

Aoki, T., S. Schweinsberg, J. Manasson and P. Schedl, 2008 A stage-specific factor confers Fab-7 boundary activity during early embryogenesis in Drosophila. Mol Cell Biol 28: 1047–1060.

Ayme-Southgate, A., J. Vigoreaux, G. Benian and M. L. Pardue, 1991 Drosophila has a twitchin/titin-related gene that appears to encode projectin. Proc Natl Acad Sci U S A 88: 7973–7977.

Bantignies, F., C. Grimaud, S. Lavrov, M. Gabut and G. Cavalli, 2003 Inheritance of Polycomb-dependent chromosomal interactions in Drosophila. Genes Dev 17: 2406–2420.

Bellen, H. J., C. J. O’Kane, C. Wilson, U. Grossniklaus, R. K. Pearson et al., 1989 P-element-mediated enhancer detection: a versatile method to study development in Drosophila. Genes Dev 3: 1288–1300.

Blanton, J., M. Gaszner and P. Schedl, 2003 Protein:protein interactions and the pairing of boundary elements in vivo. Genes Dev 17: 664–675.

Bonchuk, A., O. Maksimenko, O. Kyrchanova, T. Ivlieva, V. Mogila et al., 2015 Functional role of dimerization and CP190 interacting domains of CTCF protein in Drosophila melanogaster. BMC Biol 13: 63.

Byrd, K., and V. G. Corces, 2003 Visualization of chromatin domains created by the gypsy insulator of Drosophila. J Cell Biol 162: 565–574.

Cavalli, G., and R. Paro, 1998 The Drosophila Fab-7 chromosomal element conveys epigenetic inheritance during mitosis and meiosis. Cell 93: 505–518.

Cherbas, L., A. Willingham, D. Zhang, L. Yang, Y. Zou et al., 2011 The transcriptional diversity of 25 Drosophila cell lines. Genome Res 21: 301–314.

Chintapalli, V. R., J. Wang and J. A. Dow, 2007 Using FlyAtlas to identify better Drosophila melanogaster models of human disease. Nat Genet 39: 715–720.

Comet, I., B. Schuettengruber, T. Sexton and G. Cavalli, 2011 A chromatin insulator driving three-dimensional Polycomb response element (PRE) contacts and Polycomb association with the chromatin fiber. Proc Natl Acad Sci U S A 108: 2294–2299.

Cubenas-Potts, C., M. J. Rowley, X. Lyu, G. Li, E. P. Lei et al., 2017 Different enhancer classes in Drosophila bind distinct architectural proteins and mediate unique chromatin interactions and 3D architecture. Nucleic Acids Res 45: 1714–1730.

Czerny, T., G. Halder, U. Kloter, A. Souabni, W. J. Gehring et al., 1999 twin of eyeless, a second Pax-6 gene of Drosophila, acts upstream of eyeless in the control of eye development. Mol Cell 3: 297–307.

Dekker, J., K. Rippe, M. Dekker and N. Kleckner, 2002 Capturing chromosome conformation. Science 295: 1306–1311.

Desplan, C., 1997 Eye development: governed by a dictator or a junta? Cell 91: 861–864.

Dreos, R., G. Ambrosini, R. C. Perier and P. Bucher, 2015 The Eukaryotic Promoter Database: expansion of EPDnew and new promoter analysis tools. Nucleic Acids Res 43: D92–96.

Eagen, K. P., E. L. Aiden and R. D. Kornberg, 2017 Polycomb-mediated chromatin loops revealed by a subkilobase-resolution chromatin interaction map. Proc Natl Acad Sci U S A 114: 8764–8769.

Filion, G. J., J. G. van Bemmel, U. Braunschweig, W. Talhout, J. Kind et al., 2010 Systematic protein location mapping reveals five principal chromatin types in Drosophila cells. Cell 143: 212–224.

Frise, E., A. S. Hammonds and S. E. Celniker, 2010 Systematic image-driven analysis of the spatial Drosophila embryonic expression landscape. Mol Syst Biol 6: 345.

Fyrberg, C. C., S. Labeit, B. Bullard, K. Leonard and E. Fyrberg, 1992 Drosophila projectin: relatedness to titin and twitchin and correlation with lethal(4) 102 CDa and bent- dominant mutants. Proc Biol Sci 249: 33–40.

Gambetta, M. C., and E. E. M. Furlong, 2018 The Insulator Protein CTCF Is Required for Correct Hox Gene Expression, but Not for Embryonic Development in Drosophila. Genetics 210: 129–136.

Gaszner, M., and G. Felsenfeld, 2006 Insulators: exploiting transcriptional and epigenetic mechanisms. Nat Rev Genet 7: 703–713.

Gerasimova, T. I., and V. G. Corces, 2001 Chromatin insulators and boundaries: effects on transcription and nuclear organization. Annu Rev Genet 35: 193–208.

Geyer, P. K., C. Spana and V. G. Corces, 1986 On the molecular mechanism of gypsy-induced mutations at the yellow locus of Drosophila melanogaster. EMBO J 5: 2657–2662.

Hagege, H., P. Klous, C. Braem, E. Splinter, J. Dekker et al., 2007 Quantitative analysis of chromosome conformation capture assays (3C-qPCR). Nat Protoc 2: 1722–1733.

Hagstrom, K., M. Muller and P. Schedl, 1996 Fab-7 functions as a chromatin domain boundary to ensure proper segment specification by the Drosophila bithorax complex. Genes Dev 10: 3202–3215.

Hagstrom, K., M. Muller and P. Schedl, 1997 A Polycomb and GAGA dependent silencer adjoins the Fab-7 boundary in the Drosophila bithorax complex. Genetics 146: 1365–1380.

Halder, G., P. Callaerts, S. Flister, U. Walldorf, U. Kloter et al., 1998 Eyeless initiates the expression of both sine oculis and eyes absent during Drosophila compound eye development. Development 125: 2181–2191.

Hartig, S. M., 2013 Basic image analysis and manipulation in ImageJ. Curr Protoc Mol Biol Chapter 14: Unit14 15.

Hauck, B., W. J. Gehring and U. Walldorf, 1999 Functional analysis of an eye specific enhancer of the eyeless gene in Drosophila. Proc Natl Acad Sci U S A 96: 564–569.

Hou, C., L. Li, Z. S. Qin and V. G. Corces, 2012 Gene density, transcription, and insulators contribute to the partition of the Drosophila genome into physical domains. Mol Cell 48: 471–484.

Kassis, J. A., 1994 Unusual properties of regulatory DNA from the Drosophila engrailed gene: three “pairing-sensitive” sites within a 1.6-kb region. Genetics 136: 1025–1038.

Kassis, J. A., 2002 Pairing-sensitive silencing, polycomb group response elements, and transposon homing in Drosophila. Adv Genet 46: 421–438.

Kassis, J. A., and J. L. Brown, 2013 Polycomb group response elements in Drosophila and vertebrates. Adv Genet 81: 83–118.

Kellum, R., and P. Schedl, 1991 A position-effect assay for boundaries of higher order chromosomal domains. Cell 64: 941–950.

Lanzuolo, C., V. Roure, J. Dekker, F. Bantignies and V. Orlando, 2007 Polycomb response elements mediate the formation of chromosome higher-order structures in the bithorax complex. Nat Cell Biol 9: 1167–1174.

Li, L., X. Lyu, C. Hou, N. Takenaka, H. Q. Nguyen et al., 2015 Widespread rearrangement of 3D chromatin organization underlies polycomb-mediated stress-induced silencing. Mol Cell 58: 216–231.

Lin, N., X. Li, K. Cui, I. Chepelev, F. Tie et al., 2011 A barrier-only boundary element delimits the formation of facultative heterochromatin in Drosophila melanogaster and vertebrates. Mol Cell Biol 31: 2729–2741.

Lo, P. C., and M. Frasch, 1999 Sequence and expression of myoglianin, a novel Drosophila gene of the TGF-beta superfamily. Mech Dev 86: 171–175.

Lopez-Delisle, L., L. Rabbani, J. Wolff, V. Bhardwaj, R. Backofen et al., 2021 pyGenomeTracks: reproducible plots for multivariate genomic datasets. Bioinformatics 37: 422–423.

Maeda, R. K., and F. Karch, 2006 The ABC of the BX-C: the bithorax complex explained. Development 133: 1413–1422.

Maroto, M., J. Vinos, R. Marco and M. Cervera, 1992 Autophosphorylating protein kinase activity in titin-like arthropod projectin. J Mol Biol 224: 287–291.

Marshall, W. F., A. F. Dernburg, B. Harmon, D. A. Agard and J. W. Sedat, 1996 Specific interactions of chromatin with the nuclear envelope: positional determination within the nucleus in Drosophila melanogaster. Mol Biol Cell 7: 825–842.

Matzat, L. H., R. K. Dale, N. Moshkovich and E. P. Lei, 2012 Tissue-specific regulation of chromatin insulator function. PLoS Genet 8: e1003069.

Mihaly, J., S. Barges, L. Sipos, R. Maeda, F. Cleard et al., 2006 Dissecting the regulatory landscape of the Abd-B gene of the bithorax complex. Development 133: 2983–2993.

Mirkovitch, J., M. E. Mirault and U. K. Laemmli, 1984 Organization of the higher-order chromatin loop: specific DNA attachment sites on nuclear scaffold. Cell 39: 223–232.

Mishra, K., V. S. Chopra, A. Srinivasan and R. K. Mishra, 2003 Trl-GAGA directly interacts with lola like and both are part of the repressive complex of Polycomb group of genes. Mech Dev 120: 681–689.

Mishra, R. K., and F. Karch, 1999 Boundaries that demarcate structural and functional domains of chromatin. Journal of Biosciences 24: 377–399.

Mishra, R. K., J. Mihaly, S. Barges, A. Spierer, F. Karch et al., 2001 The iab-7 polycomb response element maps to a nucleosome-free region of chromatin and requires both GAGA and pleiohomeotic for silencing activity. Mol Cell Biol 21: 1311–1318.

mod, E. C., S. Roy, J. Ernst, P. V. Kharchenko, P. Kheradpour et al., 2010 Identification of functional elements and regulatory circuits by Drosophila modENCODE. Science 330: 1787–1797.

Naumova, N., E. M. Smith, Y. Zhan and J. Dekker, 2012 Analysis of long-range chromatin interactions using Chromosome Conformation Capture. Methods 58: 192–203.

Negre, N., C. D. Brown, P. K. Shah, P. Kheradpour, C. A. Morrison et al., 2010 A comprehensive map of insulator elements for the Drosophila genome. PLoS Genet 6: e1000814.

Ogiyama, Y., B. Schuettengruber, G. L. Papadopoulos, J. M. Chang and G. Cavalli, 2018 Polycomb-Dependent Chromatin Looping Contributes to Gene Silencing during Drosophila Development. Mol Cell 71: 73–88 e75.

Pai, C. Y., E. P. Lei, D. Ghosh and V. G. Corces, 2004 The centrosomal protein CP190 is a component of the gypsy chromatin insulator. Mol Cell 16: 737–748.

Pathak, R. U., N. Rangaraj, S. Kallappagoudar, K. Mishra and R. K. Mishra, 2007 Boundary element-associated factor 32B connects chromatin domains to the nuclear matrix. Mol Cell Biol 27: 4796–4806.

Pathak, R. U., A. Srinivasan and R. K. Mishra, 2014 Genome-wide mapping of matrix attachment regions in Drosophila melanogaster. BMC Genomics 15: 1022.

Pichaud, F., and C. Desplan, 2001 A new visualization approach for identifying mutations that affect differentiation and organization of the Drosophila ommatidia. Development 128: 815–826.

Pickersgill, H., B. Kalverda, E. de Wit, W. Talhout, M. Fornerod et al., 2006 Characterization of the Drosophila melanogaster genome at the nuclear lamina. Nat Genet 38: 1005–1014.

Quiring, R., U. Walldorf, U. Kloter and W. J. Gehring, 1994 Homology of the eyeless gene of Drosophila to the Small eye gene in mice and Aniridia in humans. Science 265: 785–789.

Ramirez, F., V. Bhardwaj, L. Arrigoni, K. C. Lam, B. A. Gruning et al., 2018 High-resolution TADs reveal DNA sequences underlying genome organization in flies. Nat Commun 9: 189.

Robinson, J. T., D. Turner, N. C. Durand, H. Thorvaldsdottir, J. P. Mesirov et al., 2018 Juicebox.js Provides a Cloud-Based Visualization System for Hi-C Data. Cell Syst 6: 256–258 e251.

Ronshaugen, M., and M. Levine, 2004 Visualization of trans-homolog enhancer-promoter interactions at the Abd-B Hox locus in the Drosophila embryo. Dev Cell 7: 925–932.

Sexton, T., F. Bantignies and G. Cavalli, 2009 Genomic interactions: chromatin loops and gene meeting points in transcriptional regulation. Semin Cell Dev Biol 20: 849–855.

Sexton, T., E. Yaffe, E. Kenigsberg, F. Bantignies, B. Leblanc et al., 2012 Three-dimensional folding and functional organization principles of the Drosophila genome. Cell 148: 458–472.

Siegal, M. L., and D. L. Hartl, 2000 Application of Cre/loxP in Drosophila. Site-specific recombination and transgene coplacement. Methods Mol Biol 136: 487–495.

Sigrist, C. J., and V. Pirrotta, 1997 Chromatin insulator elements block the silencing of a target gene by the Drosophila polycomb response element (PRE) but allow trans interactions between PREs on different chromosomes. Genetics 147: 209–221.

Srinivasan, A., and R. K. Mishra, 2012 Chromatin domain boundary element search tool for Drosophila. Nucleic Acids Res 40: 4385–4395.

Srinivasan, A., and R. K. Mishra, 2020 Genomic organization of Polycomb Response Elements and its functional implication in Drosophila and other insects. J Biosci 45.

Sultana, H., S. Verma and R. K. Mishra, 2011 A BEAF dependent chromatin domain boundary separates myoglianin and eyeless genes of Drosophila melanogaster. Nucleic Acids Res 39: 3543–3557.

Thomas, S., X. Y. Li, P. J. Sabo, R. Sandstrom, R. E. Thurman et al., 2011 Dynamic reprogramming of chromatin accessibility during Drosophila embryo development. Genome Biol 12: R43.

Valenzuela, L., and R. T. Kamakaka, 2006 Chromatin insulators. Annu Rev Genet 40: 107–138.

Vasanthi, D., A. Nagabhushan, N. K. Matharu and R. K. Mishra, 2013 A functionally conserved Polycomb response element from mouse HoxD complex responds to heterochromatin factors. Sci Rep 3: 3011.

Viets, K., M. E. G. Sauria, C. Chernoff, R. Rodriguez Viales, M. Echterling et al., 2019 Characterization of Button Loci that Promote Homologous Chromosome Pairing and Cell-Type-Specific Interchromosomal Gene Regulation. Dev Cell 51: 341–356 e347.

Vogelmann, J., A. Le Gall, S. Dejardin, F. Allemand, A. Gamot et al., 2014 Chromatin insulator factors involved in long-range DNA interactions and their role in the folding of the Drosophila genome. PLoS Genet 10: e1004544.

Wang, Q., Q. Sun, D. M. Czajkowsky and Z. Shao, 2018 Sub-kb Hi-C in D. melanogaster reveals conserved characteristics of TADs between insect and mammalian cells. Nat Commun 9: 188.

West, A. G., M. Gaszner and G. Felsenfeld, 2002 Insulators: many functions, many mechanisms. Genes Dev 16: 271–288.

Wolff, J., V. Bhardwaj, S. Nothjunge, G. Richard, G. Renschler et al., 2018 Galaxy HiCExplorer: a web server for reproducible Hi-C data analysis, quality control and visualization. Nucleic Acids Res 46: W11–W16.

Wolle, D., F. Cleard, T. Aoki, G. Deshpande, P. Schedl et al., 2015 Functional Requirements for Fab-7 Boundary Activity in the Bithorax Complex. Mol Cell Biol 35: 3739–3752.

Zhang, Y., C. H. Wong, R. Y. Birnbaum, G. Li, R. Favaro et al., 2013 Chromatin connectivity maps reveal dynamic promoter-enhancer long-range associations. Nature 504: 306–310.

